# ZEB1 Is a Key Regulator of Cardiomyocyte Structure and Function

**DOI:** 10.1101/2025.10.08.681108

**Authors:** Parul Gupta, Daniel Finke, Verena Kamuf-Schenk, Anushka Deshpande, Marc P. Stemmler, Simone Brabletz, Thomas Brabletz, Norbert Frey, Tobias Jakobi, Mirko Völkers, Vivien Kmietczyk

**Affiliations:** Department of Internal Medicine III (Cardiology, Angiology, and Pneumology), Heidelberg University Hospital, 69120 Heidelberg, Germany; DZHK (German Center for Cardiovascular Research), Partner Site Heidelberg/Mannheim, 69120 Heidelberg, Germany; Department of Experimental Medicine I, Friedrich-Alexander-University of Erlangen-Nuernberg, 91054 Erlangen, Germany; Department of Internal Medicine and the Translational Cardiovascular Research Center, University of Arizona College of Medicine - Phoenix, Phoenix, USA

**Keywords:** ZEB1, cardiac remodeling, dilated cardiomyopathy, sarcomeric integrity, heart failure, Mitochondrial defect

## Abstract

**Background:** Our previous work identified the transcription factor “Zinc Finger E-Box Binding Homeobox 1” (ZEB1) as a downstream effector of Cytoplasmic Polyadenylation Element Binding Protein 4 (CPEB4), an RNA-binding protein responsive to cardiac stress. While ZEB1 is known for its role in cancer metastasis and epithelial-to-mesenchymal transition (EMT), its function in cardiomyocytes is not well understood. Based on previous findings, we hypothesize that ZEB1 is essential for maintaining the structural integrity and mitochondrial function of cardiomyocytes.

**Methods:** AAV9-Zeb1 was used for the overexpression of Zeb1. Using a myosin heavy chain alpha (αMHC) Cre system, we created a Zeb1 conditional knockout mouse. To evaluate cardiac function and structure, we used echocardiography, electron microscopy, immunohistochemistry. We identified differentially expressed genes following Zeb1 deletion using RNA-seq and determined direct Zeb1 target genes by integrating this transcriptome data with a Zeb1 ChIP-seq dataset.

**Results:** ZEB1 deletion leads to sarcomere damage, mitochondrial dysfunction, and dedifferentiation, with more pronounced effects in females. Overexpression promotes hypertrophic remodeling. Echocardiographic analysis showed progressive systolic dysfunction, and histology revealed sarcomeric disarray, again especially in females. A tamoxifen-inducible ZEB1 knockout mouse model only confirmed ZEB1’s crucial role in fully differentiated cardiomyocytes in female mice. Integrated analysis of RNA-seq and ChIP-seq revealed that Zeb1 directly regulated mitochondrial genes, thereby playing a critical role cardiomyocyte energy supply and having secondary effects on cardiac structure and function.

**Conclusions:** ZEB1 is critical for cardiomyocyte homeostasis, and maintaining its function is necessary for normal cardiac performance and structure.

## Introduction

Cardiomyocytes are the contractile cells of the heart, originating from mesodermal progenitors and undergoing tightly regulated differentiation and maturation processes [4, 8, 21]. Alterations in cardiomyocyte homeostasis are implicated in the development of multiple cardiac diseases, such as hypertrophic and dilated cardiomyopathy, both of which play a substantial role in the incidence of heart failure and associated mortality (5–9).

The transcriptional repressor zinc finger E-box binding homeobox 1 (ZEB1) is primarily known for its function in the EMT [23, 30], which is essential for development, wound healing, and the proliferation of cancer cells [7, 30, 32, 41, 42]. To facilitate cellular plasticity and mobility, ZEB1 suppresses epithelial markers like E-cadherin (*Cdh1*) and increases the expression of mesenchymal genes like N-Cadherin (*Cdh2*) [16, 36]. In skeletal muscle, Zeb1 preserves the mesenchymal identity of satellite cells and is crucial for effective muscle repair [38]. Previous research on Zeb1 have primarily focused on its functions in cancer biology [22, 43, 45], stem cell maintenance [11], tissue fibrosis [44, 46], and regeneration [38]. Functional genomic studies in murine models have demonstrated that ZEB1 is indispensable for early mesendodermal differentiation [11, 28, 29], including cardiovascular lineage specification [10]. Knockout studies in mice show that loss of ZEB1 leads to complex developmental defects, including sporadic vascular changes and hemorrhages, indicating a role in vascular and cardiac development [25]. ZEB1, together with co-factors like HDAC2, works as a key regulator of vascular tone and cardiovascular health [10].

We previously performed a comprehensive proteome-wide screen to identify RNA-binding proteins (RBPs) and translationally regulated mRNAs associated with pathological cardiomyocyte growth. This investigation revealed that the *Zeb1* transcripts undergo post-transcriptional regulation mediated by specific RBPs. Notably, we showed that elevated Zeb1 protein expression, in the absence of corresponding changes in mRNA levels, is dependent on the activity of specific RBPs and pressure overload induced heart failure is associated with increased protein levels of Zeb1 [34]. Still the function of Zeb1 in adult cardiac myocytes and its involvement in pathological remodeling remains unexplored.

Given ZEB1’s emerging role as a regulator of mesenchymal identity, its role in cardiomyocyte differentiation and our previous findings of differential regulation in cardiomyocyte hypertrophy, we hypothesize that ZEB1 also assumes important functions in adult cardiomyocyte homeostasis and pathological remodeling.

## Materials and Methods

### Human heart failure samples

The human samples were provided by the Rangrez Lab, Internal Medicine III of the University Hospital Heidelberg. Samples had been obtained in compliance with the ethical committee of the medical school of the Georg-August-University, Göttingen, Germany[5].

### Animal Model

All animal procedures were approved by the Baden-Württemberg council (Approval: G-190/21) and conducted in accordance with animal welfare regulations.

Zeb1 overexpression was achieved by tail vein injection of 1,6*10^15 vg/animal of the AAV9 containing the murine Zeb1 coding sequence under the hTnt promoter. Equal amounts of control virus were used.

Floxed *Zeb1* mice on a C57BL/6N background were generated in the laboratory of Prof. Dr. Thomas Brabletz at the Clinical Molecular Biology Research Center (KMFZ), Friedrich-Alexander University Erlangen-Nuremberg [6]. Floxed *Zeb1* mice were crossed with the αMHC-Cre and the αMHC-MerCreMer line to generate the conditional and the tamoxifen-inducible conditional Zeb1 cKO model, respectively. Mice were bred and housed in Heidelberg’s central animal facility under controlled temperature and humidity, with a 12:12 hour light-dark cycle, and provided ad libitum access to standard chow.

Murine cardiac lysates of the RBM20 KO and the Mybpc3 KO mice were provided kindly from the van den Hoogenhof and the Backs lab (University Hospital Heidelberg, Experimental Cardiology).

### Isolation of adult mouse cardiomyocytes (AMCM)

Adult cardiomyocytes from 8-week ZEB1KO and WT mice were isolated using a Langendorff-free method [1]. Hearts were perfused with EDTA, perfusion, and collagenase buffers, then minced and dissociated. Digestion was stopped, cells filtered, calcium gradually reintroduced, and cardiomyocytes plated (∼50,000cells/cm²) on laminin-coated plates for culture

### Isolation of neonatal rat cardiomyocytes

Neonatal rat cardiomyocytes (NRCM) were isolated from 1–3-day-old Wistar rats by enzymatic digestion of ventricles. After pre-plating to remove noncardiomyocytes, cells were plated at 50,000 cells/cm² on gelatin-coated plates and maintained at 37 °C, 5% CO₂. Experiments were performed the following day.

### Adenovirus and AAV9 production and viral transduction

ZEB1 entry clones (pDONR221, NM_011546.3) were subcloned into pAd/CMV/V5-DEST via Gateway LR reaction. Constructs were transformed into E. coli, verified by Sanger sequencing, and used to produce adenoviruses in HEK293A cells. AAV9 vectors were produced via PEI transfection in 50 T175 HEK293T flasks using helper and ITR plasmids.

### Echocardiography

Echocardiography was performed using the Visual Sonics Vevo F2 system with a UHF46x transducer. B-mode and M-mode images were captured from parasternal long and short axes. Mice were anesthetized with 1.5% isoflurane on a 37°C heated plate. Heart function was analyzed at 350–500 bpm using VevoLab 5.9.0 software.

### Electron microscopy

Heart tissue was fixed in glutaraldehyde/paraformaldehyde, post-fixed with osmium tetroxide, stained with uranyl acetate, dehydrated, and embedded in Spurr resin. Ultrathin 70 nm sections were contrast-stained and imaged in the Electron Microscopy Core Facility (EMCF) at the Heidelberg University using a JEOL JEM-1400 TEM at 80 kV, 8kX–10kX magnification.

### Preparation of mouse heart lysates

Mice were sacrificed by cervical dislocation, the chest was opened, and the heart was carefully removed. Following a brief washing step in cold PBS, hearts were weighed and frozen in liquid nitrogen. Frozen heart tissue was then homogenized in 700 μL RIPA buffer, in screw-cap tubes with ceramic beads, using a Precellys tissue homogenizer (Bertin Technologies). To ensure complete lysis, tissue homogenates were passed several times through a syringe with a 25-gauge needle (Braun, 10162148) and incubated on ice for 5 min before centrifugation at 20,000 ×g for 15 min at 4 ◦C. Only the clear supernatant was retained for further analyses. Mouse heart fixation for histology

Mice were sacrificed by cervical dislocation, and the chest was promptly opened to expose the beating heart. 1 mL of 100 mM CdCl2 was carefully injected into the left ventricle to stop beating in diastole. A small incision was made in the right atrium to enable drainage, and 5 mL of heparin (diluted in PBS to 100 U/mL) was slowly perfused through the right ventricle into the left ventricle to remove excess blood. 10 mL of 10% formaldehyde (in PBS) was then perfused the same way to fix the heart tissue. Finally, the heart was excised, cleaned, and kept in 4% formaldehyde (in PBS) for 24 hours at 4◦C, before being transferred to fresh PBS for short-term storage or immediately embedded in paraffin.

### Mouse heart embedding and sectioning

After fixation, mouse hearts were dehydrated and infiltrated with paraffin using the HistoCore PEARL tissue processor (Leica). Next, hearts were embedded in paraffin blocks using the Histo-Core Arcadia H embedding station (Leica). Solid paraffin blocks were kept at-20◦C overnight before being cut at room temperature into 6 μm histological sections using a microtome (RM2145, Leica). Heart sections were transferred onto glass slides and left to dry overnight on a flattening table (HI1220, Leica) preheated to 37◦C. Tissue slides were then kept at room temperature until further use. Before histological staining was performed, heart sections were deparaffinized in xylene (3× 5 min) and rehydrated in a descending ethanol series (2× absolute EtOH, 1× 95 % EtOH, 1× 70 % EtOH, 5 min each) before 3 final washes in ddH2O (3 min each).

### Hematoxylin and eosin staining

The staining was performed using the Hematoxylin & Eosin fast staining kit as per the manufacturer’s instructions. Briefly, tissue slides were stained for 6 min in Hematoxylin, washed for 10 sec in tap water, and differentiated for 10 sec in 0.1% HCl. Next, slides were rinsed for 6 min in running tap water before they were stained for 30 sec in Eosin and finally washed for 30 sec in running tap water. Slides were then dehydrated (3 min in 95 % EtOH, 2× 3 min in absolute EtOH, and 3× 5 min in xylene), mounted with Eukitt quick-hardening mounting medium, and covered with a coverslip.

### Masson Trichrome staining

The staining was performed using Masson Trichrome Stain Kit as per the manufacturer’s instructions. Briefly, tissue slides were incubated in preheated Bouin’s Solution at 60◦C for 15 min, then washed with running tap water for 5 min and stained in Weigert’s Iron Hematoxylin Solution for 5 min. Following another washing step in running tap water, slides were quickly rinsed with ddH2O and incubated in Bieberich Scarlet-Acid Fuchsin solution for 5 min. After another rinse with ddH2O, slides were placed in working Phosphotungstic/Phosphomolybdic Acid Solution for 5 min and then directly transferred to Aniline Blue Solution for 5 min. Slides were incubated in 1% acetic acid for 2 min and rinsed with ddH2O. After dehydration series (3 min in 95% EtOH, 2× 3 min in absolute EtOH, and 3× 5 min in xylene), heart sections were mounted with Eukitt quick-hardening mounting medium and covered with a coverslip. Images of sections were taken on an Axio Vert.A1 microscope (Zeiss),

### Immunohistochemistry

After deparaffinization and rehydration of the tissue samples, antigen retrieval was performed by boiling heart tissue slides in citrate buffer in the microwave (700 W for 3 min, followed by 450 W for 12 min). Slides were cooled at 4 ◦C for 20 - 30 min, washed with ddH2O (2 × 3 min), and equilibrated in TN buffer (2 × 3 min). Blocking was achieved with TNB blocking solution for 60 min at room temperature. The antibodies and reagents were diluted in TNB blocking solution and used for staining. They were pipetted onto heart sections and incubated in a humid chamber overnight at 4◦C. The next day, slides were washed with TN buffer (3× 3 min) and if required, incubated with a secondary antibody (diluted in TNB blocking solution) for 2 hours at room temperature. The used antibodies and their dilutions are listed in section 2.4. After washing with TN buffer (3× 3 min), sections were mounted with Vectashield Antifade Mounting Medium containing 5 μg/mL DAPI. Specimens were imaged on a Leica SP8 confocal fluorescence microscope, and image analyses were carried out using ImageJ.

### WGA staining and quantification

Formalin-fixed, paraffin-embedded (FFPE) left ventricle sections (6 µm) were deparaffinized, rehydrated, and stained overnight at 4 °C with Alexa Fluor 488-conjugated Wheat Germ Agglutinin (1:50 in PBS) to label cardiomyocyte membranes. After PBS washes, sections were counterstained with DAPI (1 µg/mL), mounted with Vectashield Antifade Mounting Medium, and imaged using a Leica SP8 confocal fluorescence microscope. The cardiomyocyte cross-sectional area was quantified in transverse sections using ImageJ by tracing membrane-bound borders and analyzing ≥100 cells per heart from multiple fields.

### Transfection of NRCMs with small interfering RNA

24 hours after isolation and plating of NRCMs, cells were transfected with small interfering RNA (siRNA) to achieve a transient knockdown of Zeb1. For that purpose, control scrambled siRNA (siScr; Thermo Fisher Scientific, 4390843) or siRNA targeting Zeb1 transcript (siZeb1; Thermo Fisher Scientific, s130527) were delivered into NRCMs using HiPerfect transfection reagent according to the manufacturer’s protocol. Briefly, 5 μL of HiPerfect transfection reagent was used per well of a 12-well plate to a total volume of 500 μL per well, and the final siRNA concentration was kept at 100 nM. NRCMs were incubated with transfection mixes overnight, then the medium was replaced with fresh NRCM Treatment Medium, and cells were cultured for another 48 hours to ensure efficient knockdown.

### Treatment of NRCMs with phenylephrine

48 48 hours after transfection with siRNA, NRCMs were treated with alpha-1 adrenergic receptor agonist phenylephrine (PE), which simulates pathological hypertrophy by inducing NRCM growth. A fresh stock solution of PE (1 mg/mL in PBS) was prepared before each experiment and diluted in NRCM Treatment Medium to a final concentration of 50 μM. NRCMs were kept in PE-containing medium for 24 hours.

### Adenovirus production and viral transduction

Entry clones were ordered from BioCat and cloned in pDONR221 backbone with the following coding DNA sequences (CDS): pEntr_ZEB1 (NM_011546.3, M. musculus CDS). These were subcloned into pAd/CMV/V5-DEST expression vector (Invitrogen, V49320) by performing the LR reaction using Gateway LR Clonase II Enzyme mix per manufacturer’s instructions. The obtained construct was transformed into One Shot TOP10 Chemically Competent E. coli cells (Thermo Fisher Scientific, C404010) as indicated in the product information sheet, spread onto plates with LB-agar and ampicillin (100 μg/mL) and incubated at 37◦C overnight. The next day, colonies were picked and plasmids were isolated using the peqGOLD Plasmid Miniprep Kit I. The correct insert sequence was confirmed by Sanger sequencing (Mix2Seq Service, Eurofins). Glycerol stock of initial bacterial transformation was then used to prepare a liquid culture, and the expression plasmid was isolated using HiSpeed Plasmid Midi Kit. 6 μg of each expression plasmid was linearized with restriction enzyme PacI for 3 hours at 37◦C, before the enzyme was inactivated for 20 min at 65◦C. The linearized plasmid was transfected into HEK293A cells (10 cm plate) using PEI transfection reagent. Two days later, the cells were split into three 10 cm plates and cultured in antibiotic-free medium. Plates were inspected every day under a microscope and were harvested once 30 % of the cells detached. Following 3 freeze-thaw cycles, the lysate was used to reinfect HEK293A cells (T175 flask), which were harvested once 70% of the cells detached. After 3 freeze-thaw cycles, benzonase was added to the lysate, and tubes were incubated for 90 min in a heated water bath (37◦C). Finally, tubes were centrifuged at 3700 rpm for 40 min at 4◦C, the supernatant was collected and aliquoted for further use. To determine the number of adenoviral particles, a small aliquot of each virus was diluted in 0.1% SDS, and absorbance at 260 nm was measured on a NanoDrop Lite spectrophotometer (Thermo Fisher Scientific). The number of viral particles per mL was calculated as follows: A260 × dilution factor × 1.1 × 1012 particles.

### Cell lysate preparation

Cultured cells were washed with PBS and then harvested in RIPA buffer with added protease and phosphatase inhibitors. Cell lysates were collected by carefully scraping cell culture dishes and subsequently centrifuged at 20,000 x g for 15 min at 4◦C. Only soluble fractions (supernatants) were retained for further use.

### Western blotting

Protein concentration was measured by Bio-Rad DC assay after RIPA lysis. Samples were mixed with Laemmli buffer, heated, separated by SDS-PAGE, transferred to PVDF membranes, blocked, incubated with primary and secondary antibodies, and visualized using ECL detection.

**Table. 1.**
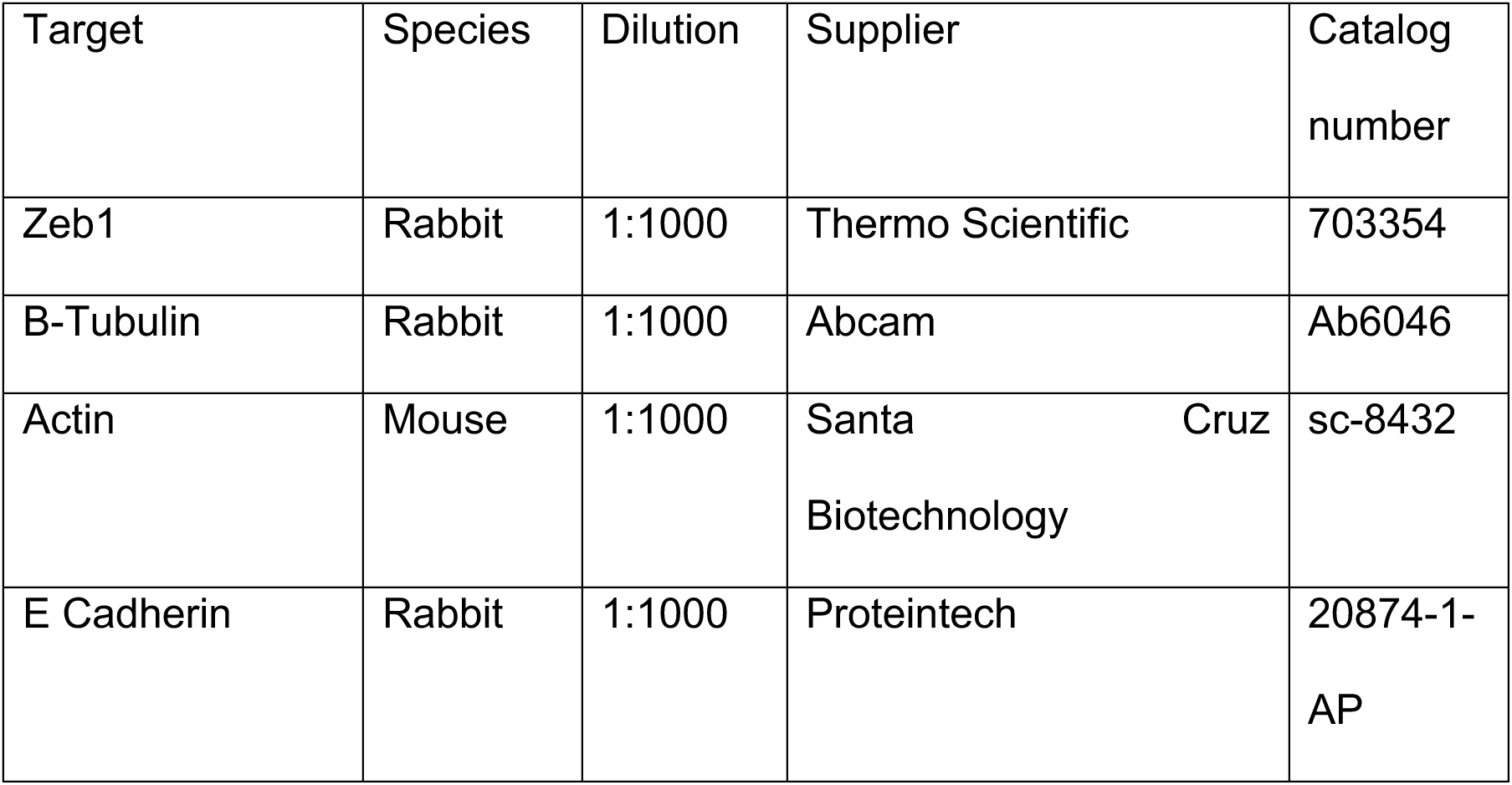
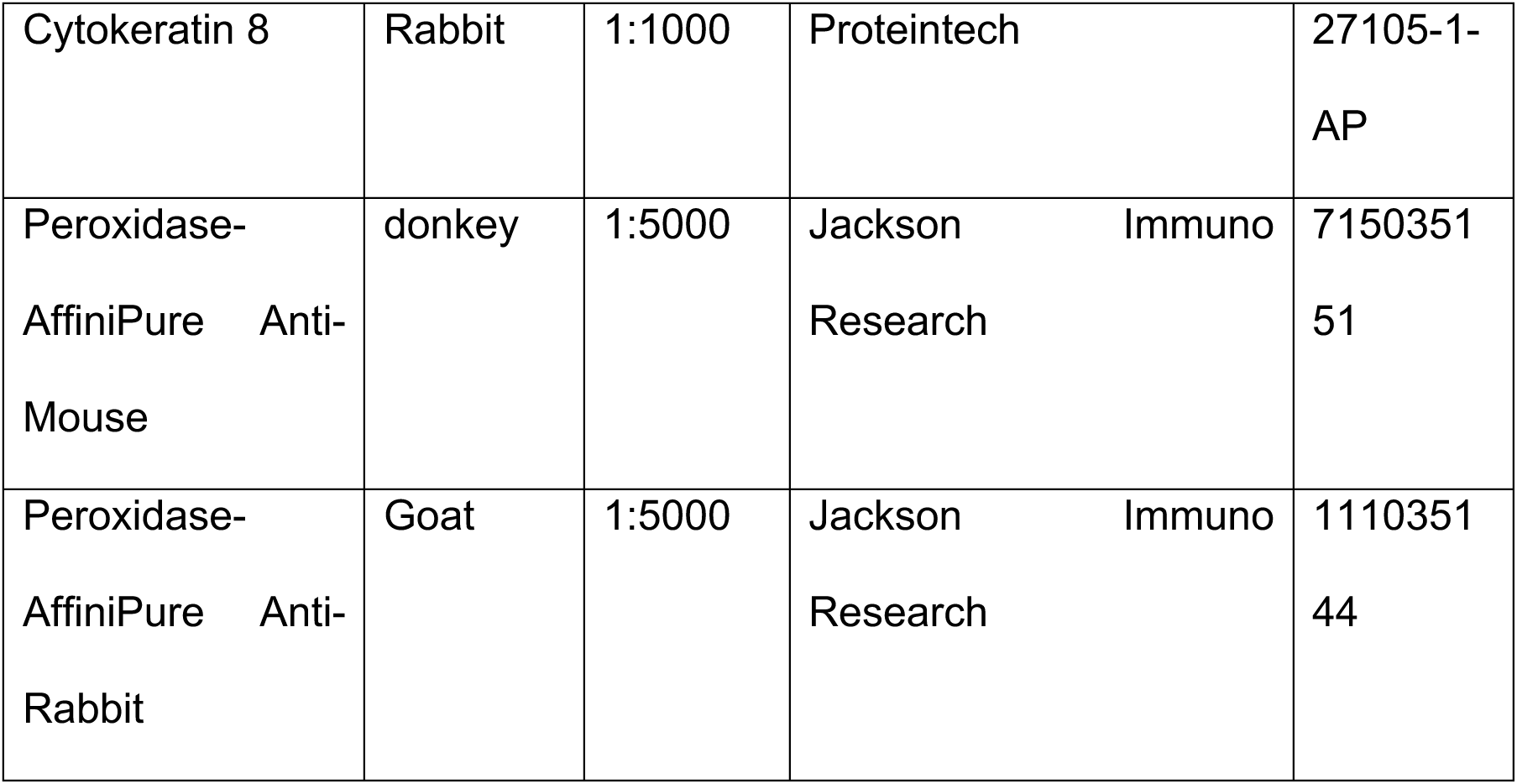
Antibodies used in western blot.

### RNA isolation and quantitative RT-PCR

Total RNA from NRCMs and homogenized. Snap-frozen tissues in RIPA buffer were isolated using Qiazol with chloroform/isopropanol precipitation. cDNA was synthesized using the iScript Kit, and qRT-PCR was performed with SYBR Green and 3μM primers. Relative expression was calculated using the ΔΔCT method with 18s or HPRT.

**Table. 2.**
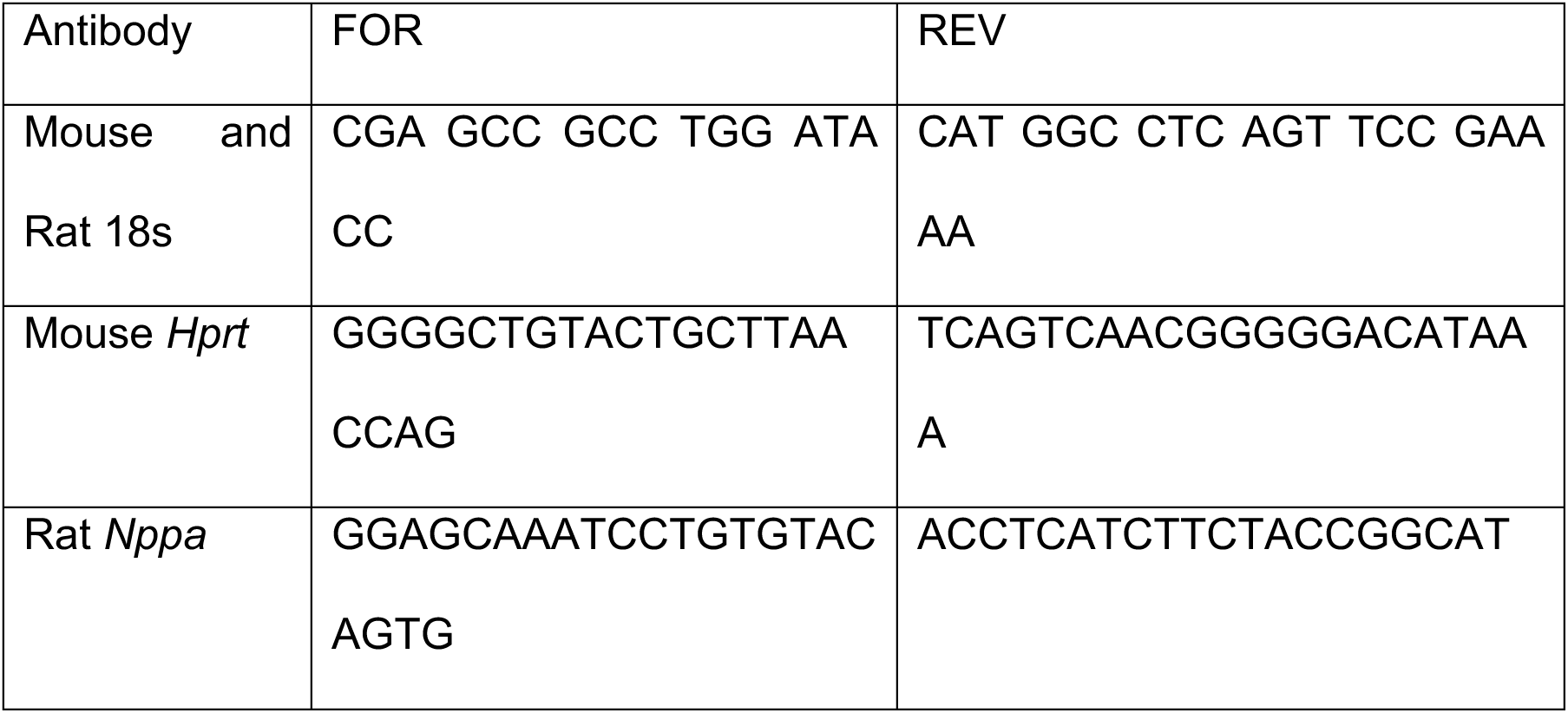
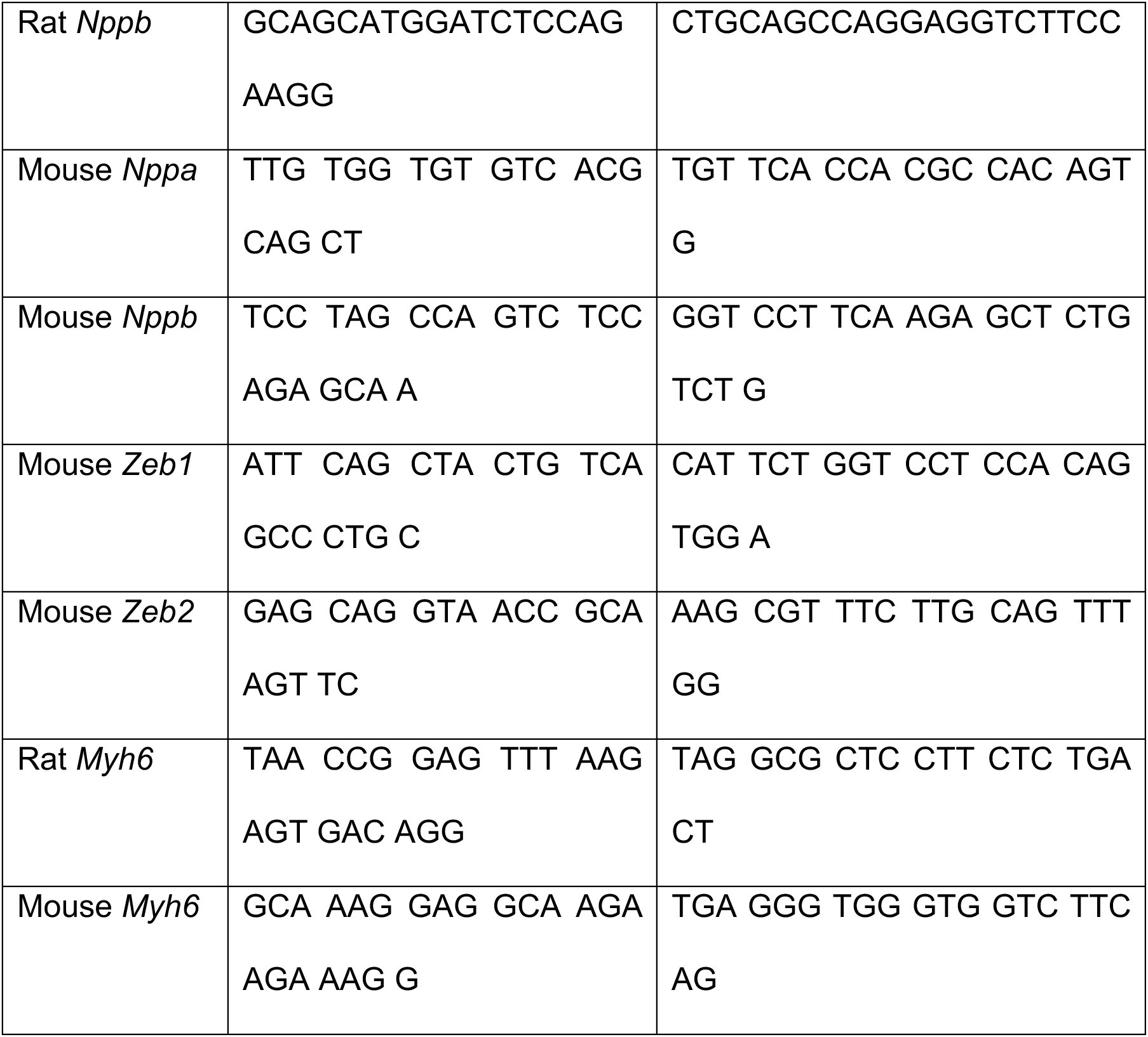
List of primers.

### Reverse transcription and quantitative real-time PCR

500 – 1000 ng of total RNA was subjected to reverse transcription with the First Strand cDNA Synthesis Kit, and oligo-dT primers were used to specifically detect mature polyadenylated mRNAs and produce a more consistent qRT-PCR signal. The resulting cDNA samples were diluted ten-to twenty-fold with nuclease-free H2O and used for quantitative real-time PCR (qRT-PCR) analysis with iTaq universal SYBR Green Supermix. Each reaction consisted of 5 μL iTaq universal SYBR Green Supermix, 300 nM forward primer, 300 nM reverse primer, and 3 μL diluted cDNA. Samples were pipetted into 96- or 384-well microplates and analyzed with the QuantStudio5 real-time PCR System (Thermo Fisher Scientific). The sequences of primers used are given in section 2.5. Each primer pair was tested before use to ensure an amplification efficiency between 90% and 110%. Additionally, melting curves were carefully inspected, and agarose gel electrophoresis was performed to verify that a single gene-specific amplicon of the correct size was produced. Ct values of genes of interest were normalized to one or the mean of two reference genes (18S rRNA and Hprt). Relative abundances and changes in expression were calculated using the 2–ΔΔCt method.

### Immunofluorescence and staining

Immunofluorescence was performed on NRCMs to assess cell size. Cells were fixed with 4% PFA, permeabilized, and blocked with 10% horse serum. Primary antibodies were applied overnight at 4°C, followed by secondary antibodies and DAPI staining. Images were captured using a Leica SP8 confocal microscope.

**Table. 3.** Antibodies used for immunofluorescence.

| Target | Species | Dilution | Supplier | Catalog number |
| --- | --- | --- | --- | --- |
| Zeb1 | Rabbit | 1:100 | Thermo Scientific | 703354 |
| Sarcomeric Actinin | Mouse | 1:1000 | Santa Cruz Biotechnology | sc-8432 |
| CSRP3 | Rabbit | 1:1000 | Proteintech | 10721-1-AP |
| Wheat Germ<br>Agglutinin, FITC<br>conjugate |  | 1:50 | Sigma-Aldrich | L4895-2MG |
| Anti-rabbit IgG-FITC | Donkey | 1:100 | Jackson Immuno<br>Research | 711-095-152 |
| Anti-rabbit IgG-Cy3 | Donkey | 1:100 | Jackson Immuno<br>Research | 711-165-152 |
| Anti-Mouse IgG-FITC | Donkey | 1:100 | Jackson Immuno<br>Research | 715-095-151 |
| Anti-Mouse IgG-Cy3 | Donkey | 1:100 | Jackson Immuno<br>Research | 715-165-151 |

### Cell size measurement

ImageJ/Fiji software (https://imagej.net/software/fiji/) was used to assess cell size. Cell size was determined manually. Using a blinded technique, in which the treatment allocation for each sample was not known. No identifying information was included in the imported photos, and each sample was given an anonymous label. An unbiased analysis was ensured by using randomized measurements to prevent biases and by compiling and analyzing the results without disclosing the treatment allocation. In each experiment, at least 100 cardiomyocytes were examined

### Bulk RNA seq

Total RNA from left ventricular tissue was extracted using TRIzol and RIPA homogenization. RNA quality (RIN>7) was confirmed via NanoDrop and Bioanalyzer. mRNA libraries were prepared with NEBNext Ultra II and sequenced (150bp paired-end) on Illumina NovaSeq. Paired-end, rRNA-depleted sequencing data were analyzed following quality control. Low-quality regions and adapter sequences were removed using Flexbar [35], and residual rRNA reads were filtered with Bowtie2 [24] using an rRNA-specific index. Reads were aligned to the ENSEMBL mouse genome build 107 (mm39) using STAR [13], and gene-level counts were obtained with Rsubread [26]. Quality metrics and mapping summaries were assessed via MultiQC [14]. Differential expression was analyzed with edgeR [12]. GO analysis was performed with topGO, using genes with RPKM≥1 across all samples as background. Heatmaps were generated using Complex Heatmap [18]. All downstream analyses were conducted in R. Sequencing raw data is available under accession number SRP630684 in the SRA, processed gene counts are available in GEO under accession number E-MTAB-15712.

### Chromatin immunoprecipitation Sequencing

NRCMs were crosslinked (40 million cells, 1% formaldehyde for 10min), quenched (250mM glycine), washed with PBS, and the nuclei were isolated. Chromatin was sonicated (3×15 cycles, Bioruptor) and cleared as described earlier [15]. ChIP was performed using ZEB1 antibody, we used 4µg of the antibody for the pulldown.

ChIP samples were treated with RNase A, Proteinase K, and SDS overnight. DNA was purified (phenol/chloroform), libraries prepared according to the manufactures’ protocols (NEBNext Ultra II) and quality-checked on TapeStation. Sequencing was performed on a DNBSEQ-G400 platform.

FASTQ data were annotated to the rat genome (rn7) using bowtie2 (version2.5.2). Only uniquely mappable reads have been subjected to further analysis. The resulting BAM files have been used for MACS (Model based analysis of ChIP-seq) peak calling (Version2.2.6) with a selected q-value (false discovery rate) of 0.05 and normalized to the corresponding input file. The DeNovo transcription factor analysis of called peaks with promoter peak was done in HOMER (version5.1) using the size given option and randomly selected background sequences.

For illustrations, bigwig files were generated via bamCoverage (version3.5.5) in Reads Per Kilobase per Million (RPKM) and 50 bp binning and averaged using bigwigAverage (version3.5.5).

The heatmaps shown were built with deeptools (version3.5.4) via usegalaxy.org [33] using a bed file of either all differentially regulated transcripts from the RNA-seq or a bed file of all summarized peaks from the single replicates of the ChIP-seq. The enrichment shown is averaged from the three replicates. Single exemplary ChIP-seq tracks of Input and pulldown are shown in Suppl.Figure.7. Peak annotation was done with HOMER (version5.1). Promoter peaks were defined as-500bps to +100bps relatively to the transcriptional start site (TSS).

The number of reads, the percentage of uniquely mappable reads and the amount of called peaks are listed separately. Raw and annotated data have been submitted to the EMBL-EBI database (accession number: E-MTAB-15712).

**Table. 4.**
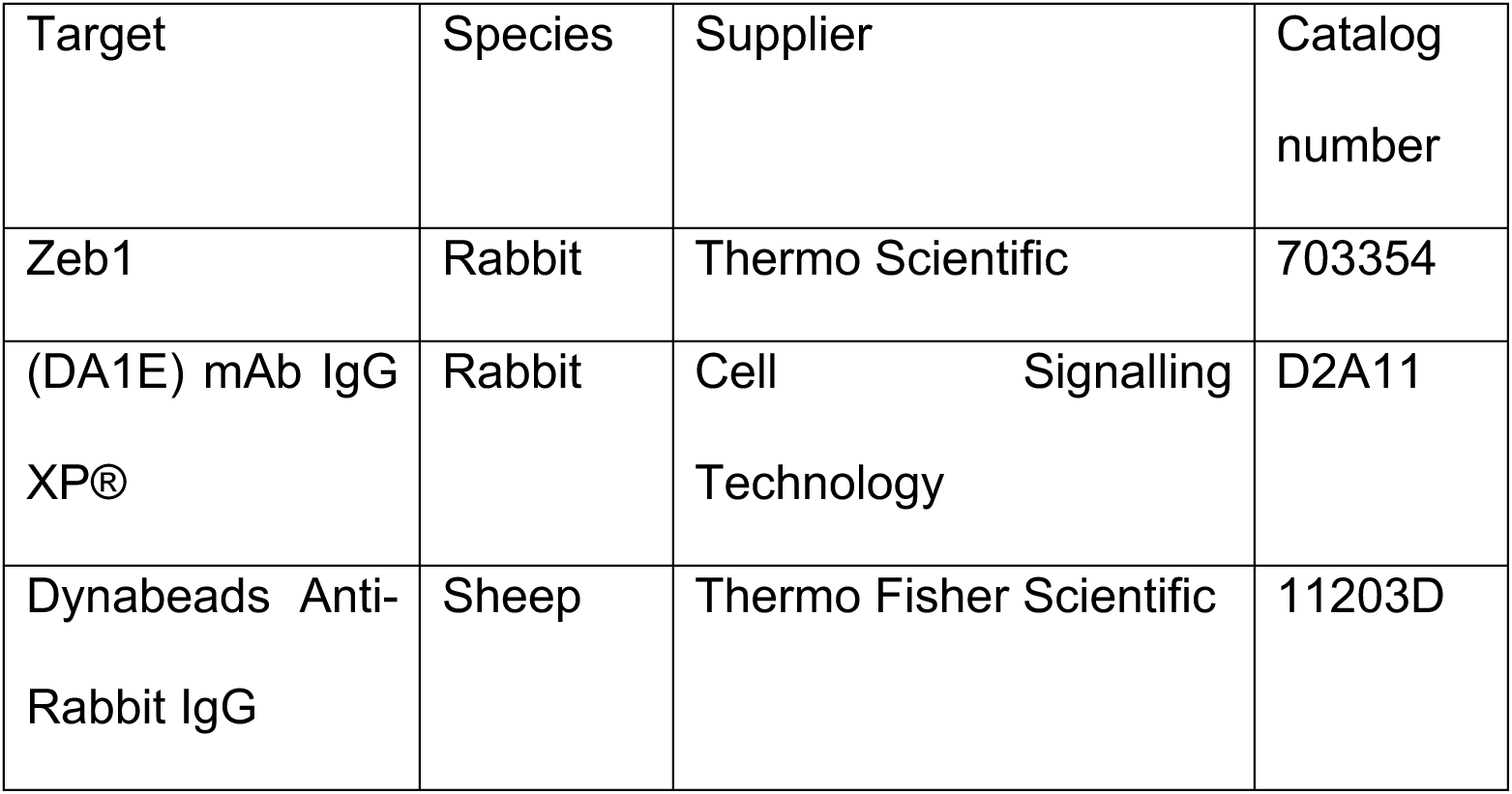
Antibodies used for Immunoprecipitation.

### Statistics and WB loading controls

Data are shown as mean ± SEM. Statistical analysis was performed using unpaired Student’s *t*-test or one-way ANOVA (*=p<0.05, **=p<0.01, ***=p<0.001, ****=p<0.0001). Tubulin, Actin, Ponceau, and Total protein stain are shown as loading controls. Data are representative of at least three biological replicates.

## Results

### Regulation of ZEB1 protein expression in cardiomyopathy models

Proteome-and transcriptome wide screening identified Zeb1 mRNA as a target of RBP-mediated post-transcriptional control in cardiomyocytes, with elevated protein levels during pressure overload–induced heart failure without corresponding mRNA changes [34]. Despite this regulation, the functional significance of Zeb1 in adult cardiac myocytes is still uncharacterized.

To test the hypothesis that dysregulation of Zeb1 expression is a common response in the heart to pathological stress, the expression of Zeb1 in different mouse models of heart failure as well as in human heart failure samples (36) was checked at protein levels. Western blotting of lysates from human cardiomyopathies and non-failing heart biopsies [5] revealed decreased Zeb1 levels (Fig.1a). Quantification of the band by densitometry (area marked by red box) and normalization to total protein stain by ponceau showed a significant reduction in ZEB1 protein levels in heart failure as compared to non-failing (NF) heart samples (Fig.1b).

**Figure 1.**
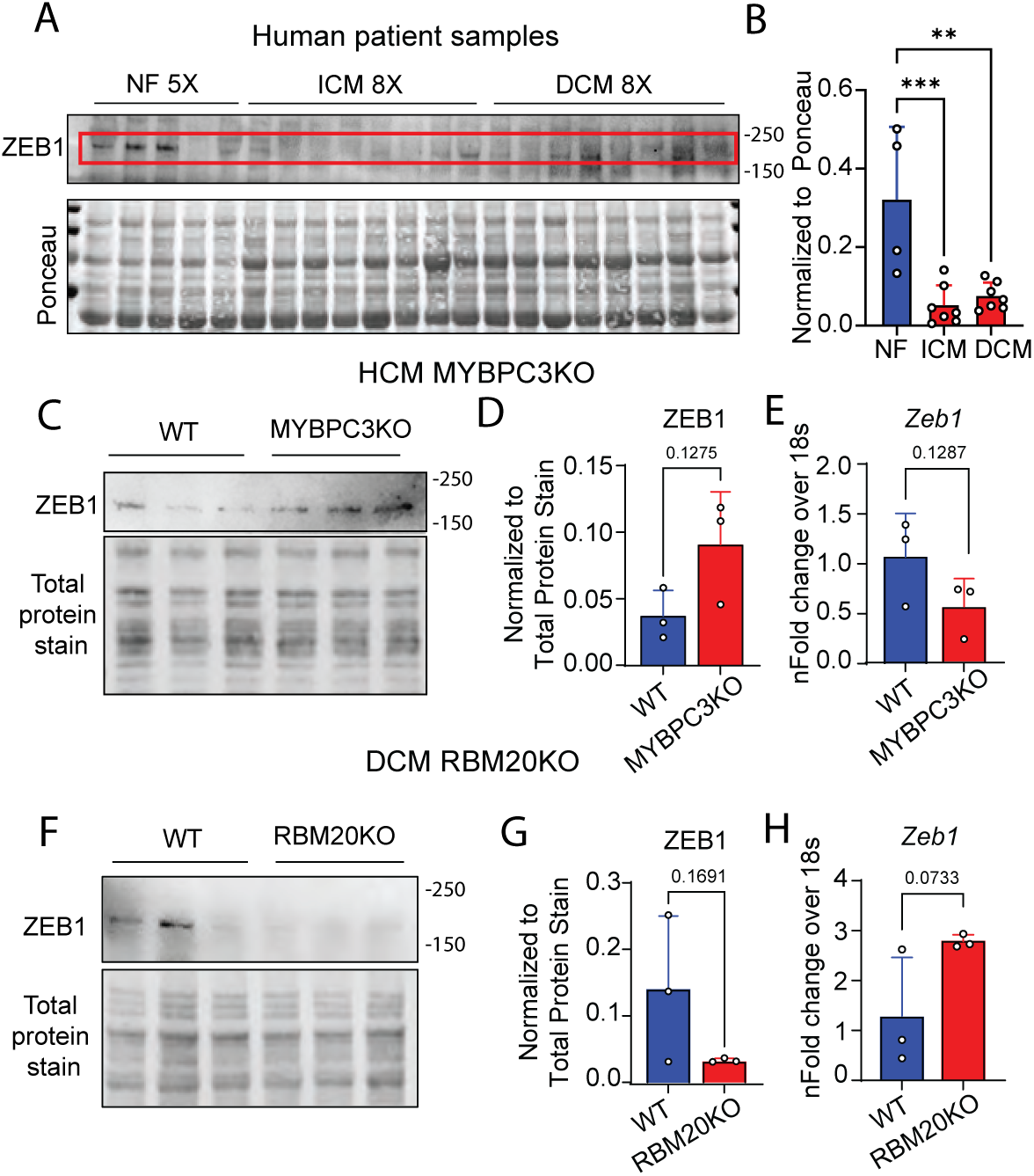
Regulation of ZEB1 protein expression in cardiomyopathy models. (a) Western Blot for Zeb1 and total protein stain by Ponceau and (b) its densiometric quantification of human ICM and DCM heart failure biopsy samples together with non-failing (NF) control samples show significantly reduced ZEB1 protein levels in ICM (n=8) and DCM (n=8) compared to NF (n=5) hearts. (c) Western Blot for Zeb1 and total protein stain of cardiac lysates of Mybpc3 knockout and control mice and (d) quantification of the immunoblot show a tendency of increased ZEB1 protein expression in the Mybpc3 knockout mice, a model of HCM (n=3/group). (e) Zeb1 mRNA expression in Mybpc3 knockout, quantified by RT-qPCR, is not significantly changed in comparison to wild type mice. (f) Western Blot for Zeb1 and total protein stain of left ventricular lysates of Rbm20 knockout mice, exhibiting DCM, and (g) quantification of the immunoblot show a tendency of decreased ZEB1 protein expression in left ventricular tissue from Rbm20 knockout mice (n=3/group). (h) Zeb1 mRNA expression in left ventricular tissue of Rbm20 knockout mjice, quantified by RT-qPCR, shows a tendency of upregulation but is not significantly changed in comparison to wild type mice. (Dilated cardiomyopathy: DCM, Ischemic cardiomyopathy: ICM, Knockout: KO, Myosin binding protein C3: Mybpc3, Non-failing: NF, Ribosome binding protein 20: Rbm20, Zinc finger E-box binding homeobox 1: Zeb1).

Western blotting of murine left ventricular lysates from Mybpc3 knockout mice, a mouse model of genetic hypertrophic cardiomyopathy (HCM) [20], revealed elevated Zeb1 protein levels (Fig.1c). Quantification of the band and normalization to total protein stain validates this tendency in comparison to wild type (WT) with a p value of 0.1275 (Fig.1d), while the transcript levels of Zeb1 remain unchanged compared to WT mice (Fig.1e). In turn, the protein levels of Zeb1 are decreased in RBM20 knockout (KO) mice, which exhibit dilated cardiomyopathy (DCM) [19], shown by western blotting (Fig. 2f) and its densitometric quantification (Fig.1g). Transcript levels of Zeb1 are not significantly changed and rather tend to be increased in the RBM20 KO (p = 0.0733).

**Figure. 2.**
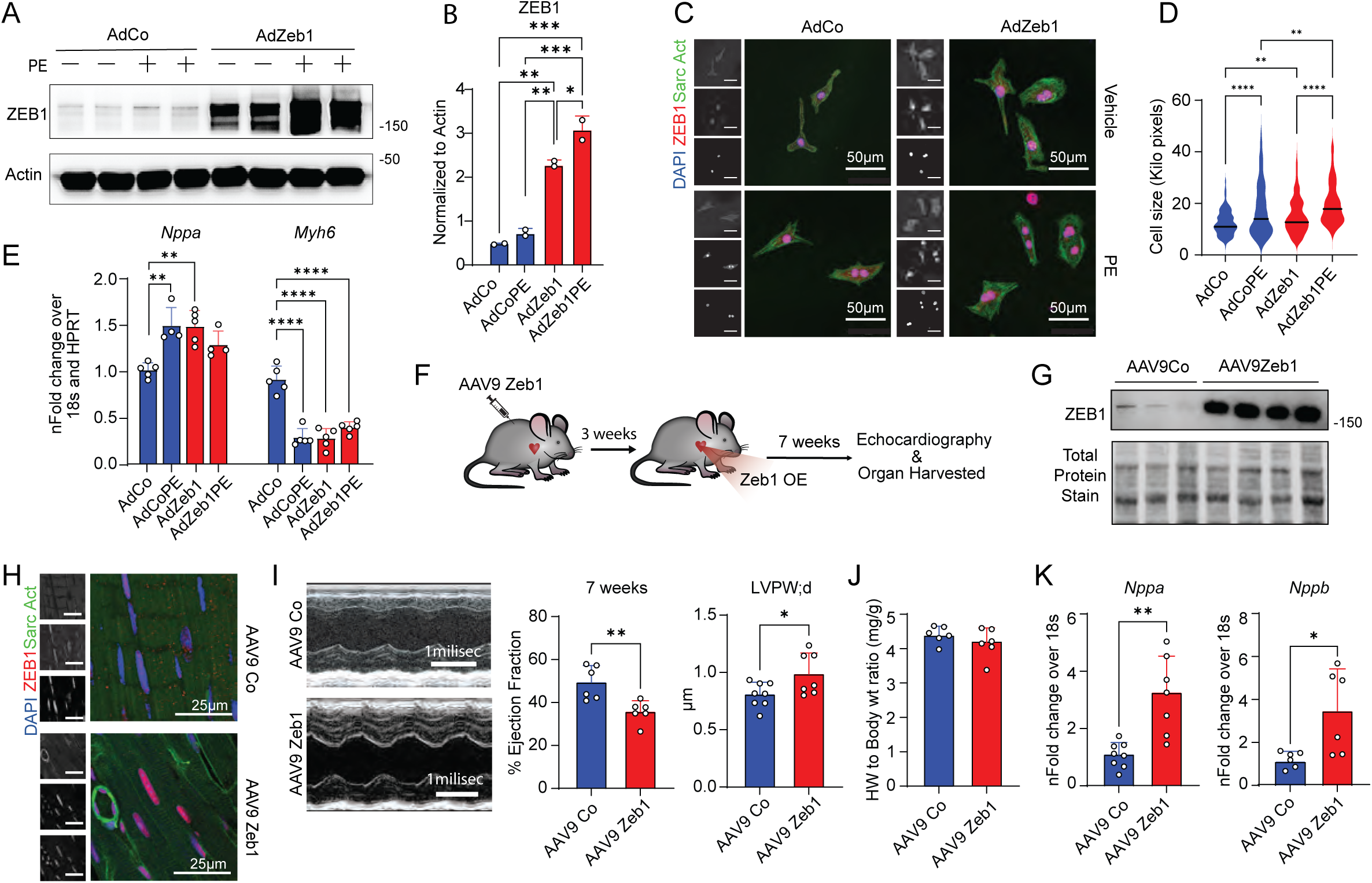
Zeb1 overexpression promotes cardiomyocyte growth *in vitro* and cardiac remodeling in male mice. (a) Western Blot for Zeb1 and beta-Actin of lysates from NRCMs transduced with adenovirus (Ad-) containing a control sequence (-Co) or coding sequence for Zeb1, treated with/without PE. (b) Quantification of the Zeb1 band from (a) showing increased Zeb1 mRNA levels in AdZeb1 transduced NRCMs. (c) Representative images of immunofluorescence staining of NRCMs (sarcomeric α-actinin - green, Zeb1 - red, and DAPI - blue) transduced with adenovirus (Ad-) containing a control sequence (-Co) or coding sequence for Zeb1, treated with/without PE. (d) Quantification of NRCM cell surface area from (c) (n = 3 independent experiments with in total > 200 cells/group). (e) *Nppa* and *Myh6* mRNA expression (n=5/group) in NRCMs transduced with adenovirus (Ad-) containing a control sequence (-Co) or coding sequence for Zeb1, treated with/without PE, quantified by RT-qPCR. (f) Schematic showing AAV9-mediated, cardiomyocyte-specific Zeb1 overexpression in adult mice and time lone of echocardiographic assessment until euthanization. (g) Western blot of left ventricular lysates of mice with AAV9-mediated, cardiomyocyte-specific Zeb1 overexpression or controls for Zeb1 and total protein stain, confirming Zeb1 upregulation in the left ventricle tissue lysates of control and Zeb1OE mice (n=4/group). (h) Representative images of immunohistochemistry (sarcomeric α-actinin-green, Zeb1-red, and DAPI-blue) confirming the cardiomyocyte specific upregulation of Zeb1. (i) Echocardiography showing representative M-mode image of the left ventricle contractions and graphical representation of significantly reduced ejection fraction, increased LVPW;d (n=6-8/group). (j) Heart/body weight ratio of mice with AAV9-mediated, cardiomyocyte-specific Zeb1 overexpression or controls (n=6/group). (k) *Nppa/Nppb* mRNA expression in mice with AAV9-mediated, cardiomyocyte-specific Zeb1 OE, quantified by RT-qPCR (n=6-8/group). (Control: Co, Left ventricular posterior wall thickness in diastole: LVPW;d, Phenylephrine: PE, Zeb1 overexpression: Zeb1OE).

In summary these data indicate dysregulation of Zeb1 in human failing hearts and in two genetic models of cardiomyopathies independent of their etiology.

### Zeb1 overexpression induces cardiomyocyte hypertrophy and cardiac dysfunction *in vitro* and *in vivo*

To determine if increased Zeb1 protein levels are associated with pathological remodeling or if Zeb1 is sufficient and necessary for pathological cell growth, we first used an adenovirus-mediated overexpression approach in neonatal rat cardiomyocytes (NRCMs). Increased ZEB1 levels induced by adenoviral transduction were confirmed by western blot and qRT-PCR (p<0.0001) (Fig.2a,b). To determine how Zeb1 overexpression affects the hypertrophic response, the α1-adrenergic receptor agonist phenylephrine (PE) was administered to both the control and Zeb1-overexpressing (Zeb1OE) NRCMs. PE treatment did increase Zeb1 protein levels in control cells without increasing transcript levels (Fig.1a,b), confirming earlier results [34].

Cell size was assessed by immunofluorescence of NRCMs overexpressing Zeb1, with or without PE treatment (Fig. 2c). The quantification of the cell area showed a significant increase (p<0.002) in cell size after Zeb1OE, similar to PE treated control cells (Fig.2d). Semi-quantitative qPCR analysis showed that NRCMs overexpressing Zeb1 displayed increased (p=0.001) expression of the hypertrophic markers *Nppa* (and *Nppb*, Supp.Fig.2a) in combination with reduction (p<0.0001) of *Myh6*, comparable to PE-treated cells (Fig.2e). In summary, overexpression of Zeb1 induces hypertrophic growth accompanied with typical increased expression of genes from the fetal gene program in NRCMs.

To validate the functional effect of Zeb1 *in vivo,* we used an Adeno-Associated Virus 9 (AAV9) cardiomyocyte-specific Zeb1 vector. Recombinant AAV9 particles were created, purified, and systemically injected into the tail veins of 8-week-old male (Fig.2f) and female mice. Increased Zeb1 expression was validated by western blot (Fig.2g), qRT-PCR (p<0.0001) (Supp.Fig.2b), and immunostaining of left ventricle tissue sections (Fig.2h).

Echocardiographic assessment up to seven weeks after injection of the AAV9 was performed to assess cardiac function. Seven weeks after the AAV9 injection, Zeb1OE male mice showed a significant decrease in ejection fraction (p=0.001), while the left ventricular posterior wall thickness during diastole (LVPW;d) was slightly but significantly increased (p=0.036), as shown in Fig. 2i. In contrast, in female mice overexpression of Zeb1 for 7 weeks did not affect ejection fraction nor LVPW;d (Supp.Fig.2c,d).

Upon Zeb1OE the heart weight to body weight ratio remained unchanged in male (Fig.2j) and female mice (Supp.Fig.2e), while the cardiomyocyte cross-sectional area, as quantified by WGA staining, showed increased cardiomyocyte size (p<0.0001) in male mice (Supp.Fig.2f,g). Additionally, expression of typical hypertrophic markers such as *Nppa* (p=0.001) and *Nppb* (p=0.02) were significantly increased in the left ventricle tissue lysates of Zeb1OE hearts as compared to control hearts, both in male and in female mice (Fig.2k and Supp.2h) suggesting pathological remodeling in male mice and to some extent female mice.

In summary, *in vivo* Zeb1 gain-of-function leads to cardiac dysfunction in male mice and the beginning of hypertrophic remodeling.

### Cardiomyocyte-specific deletion of Zeb1 leads to progressive cardiac dysfunction and pathological remodeling

To further characterize the role of Zeb1 in cardiomyocytes, we aimed to study the consequences of cardiomyocyte-specific Zeb1 depletion *in vivo*. For this, we used the αMHC-Cre mouse model to create a cardiomyocyte-specific Zeb1 conditional knockout (cKO) mouse model [6] (Fig.3a). Cardiomyocyte-specific Zeb1 deletion was validated by western blot (p=0.004) and immunostaining of left ventricular tissue, comparing the Zeb1 expression in left ventricular tissue and liver lysates of Zeb1cKO and WT adult mice (Fig.3a,b, Supp.Fig. 3a). Semi-quantitative qPCR analysis additionally confirms the KO (Supp.Fig.3B), however in both methods, western blotting and RT-qPCR Zeb1 signals are not fully depleted due to the high expression of Zeb1 in other cell types of the heart. The expression of Zeb2, Zeb1’s closest homolog, is not affected by the absence of Zeb1, both in young and old mice (Supp.Fig.3b), indicating that Zeb1 plays a non-redundant role and its function is not compensated by its homolog Zeb2 in cardiomyocytes.

**Figure. 3.**
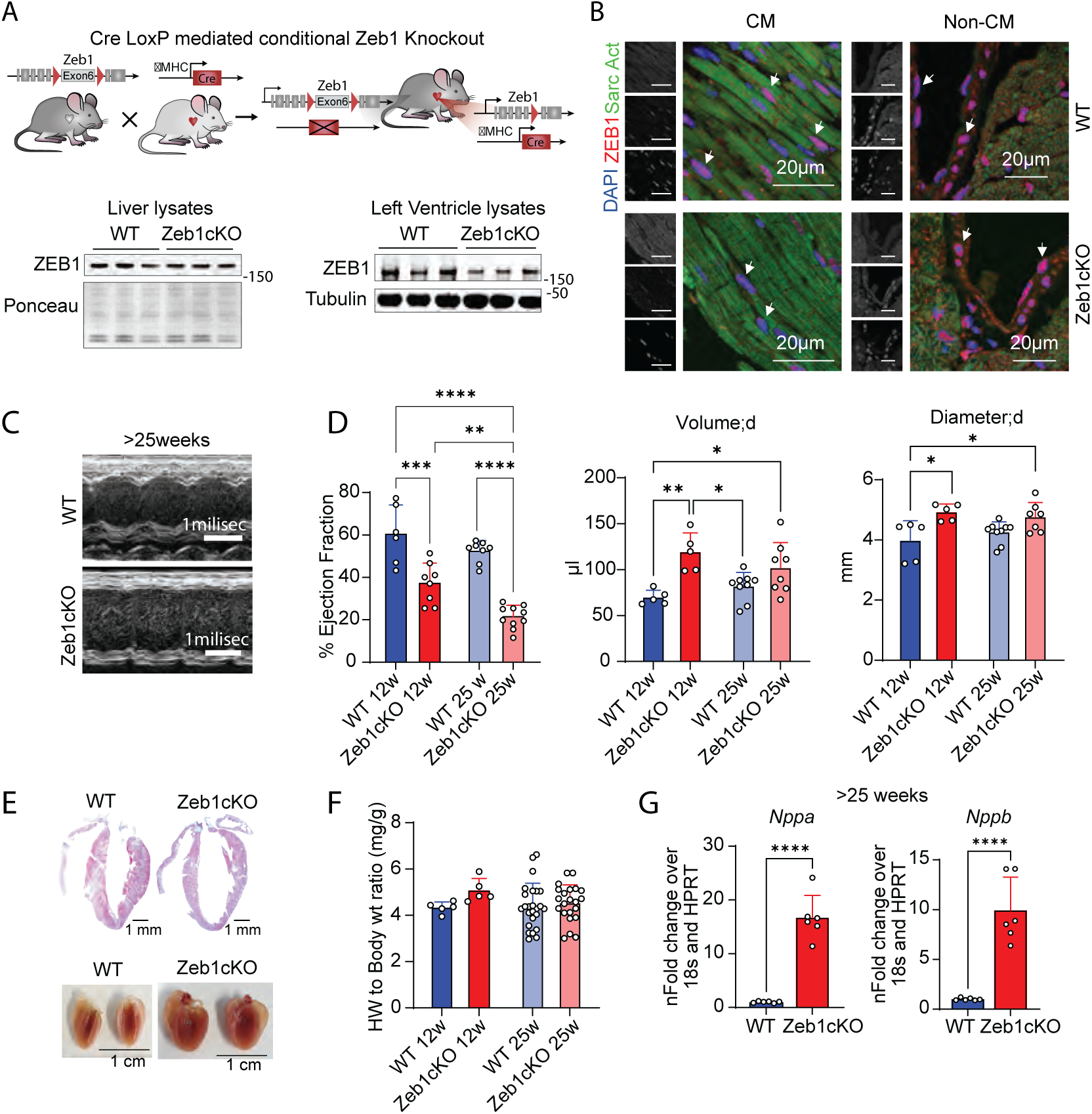
Cardiomyocyte-specific Zeb1 deletion leads to progressive cardiac dysfunction and remodeling in male mice. (a) Breeding schematic of the generation of the Zeb1cKO mice using the Cre-loxP system (Upper panel). Validation of cKO via western blot of lysates from liver and hearts of control and Zeb1cKO mice, showing Zeb1 loss in the heart but not liver (lower panel). (b) Representative images of immunostaining (sarcomeric α-actinin-green, Zeb1-red, and DAPI-blue) of left ventricular tissue sections, showing Zeb1 loss only in nuclei (white arrows) of cardiomyocytes. (c) Echocardiography showing representative M-mode image of the left ventricle contractions of 25 week and older male mice. (d) Echocardiographic analysis showing significantly reduced EF and increased LV volume and diameter in 12-week-old Zeb1cKO mice, with further EF decline by age of 25 weeks (n=5-9/group). (e) Gross morphology and histology showing ventricular dilation of 25 week and older male Zeb1cKO mice hearts fixed in formaldehyde (lower panel) and after Masson Trichrome stainings (upper panel). (f) Heart/body weight ratio of Zeb1cKO male mice are unchanged in comparison to control mice, independent of the age group shown (n=5-22/group). (g) qRT-PCR showing significantly elevated *Nppa/Nppb* mRNA levels in 25 weeks and older Zeb1cKO male left ventricle lysates in comparison to wild type animals (n=6/group). (Cardiomyocytes: CM, Cardiomyocyte-specific Zeb1 conditional knockout: Zeb1cKO, Ejection fraction: EF, Left ventricle: LV, Wild type: WT).

Echocardiography was performed to assess cardiac function and cardiac parameters of 12-week-old and 25 weeks and older mice. As early as 12 weeks of age, Zeb1cKO mice showed a significant decrease (p=0.0006) in systolic heart function, with an average Ejection Fraction (EF) of about 40% compared to 60% in WT mice (Fig.3c,d). The diastolic LV volume (p=0.003) and LV diameter (p=0.015) were significantly increased in 12-week-old male mice (Fig.3d). In older Zeb1cKO mice (≥25 weeks), the condition worsened, with the EF of Zeb1cKO mice decreasing to 20% (p<0.0001) (Fig.3d).

Accordingly, histological analysis confirmed wall thinning and dilation of the left ventricle of Zeb1cKO mice (Fig.3e), while overall heart weight to body weight ratio was not altered (Fig.3f). Dilative cardiac remodeling is characterized by pathological re-expression of embryonic genes like *Nppa* and *Nppb*. Pathological remodeling in the adult Zeb1cKO was assessed by qRT-PCR of *Nppa* (p<0.0001) and *Nppb* (p<0.0001), which revealed higher transcriptional levels in Zeb1cKO LV lysates compared to WT (Fig.3g).

### Zeb1cKO mice develop a dilated cardiomyopathy phenotype and adult-onset Zeb1 deletion causes cardiac dysfunction

Female Zeb1cKO mice exhibited a more pronounced cardiac phenotype compared to males. By 20 weeks of age, mortality began to increase among female Zeb1cKO mice, with only 80% surviving the age of 30 weeks (Fig.4a), whereas 95% of male cKO mice reached the same age (Supp.Fig.3c). Hearts of female Zeb1cKO also had severely dilated ventricles assessed by echocardiography (Fig.4b). Severe remodeling was also reflected by echocardiographic measurement showing a significant decrease in EF down to 8–10% (p<0.0001), and a significant increase in the ventricular volume (p=0.02) and diameter (p=0.031) of Zeb1cKO 25 weeks and older females (Fig.4c). The heart weight to body weight ratio was slightly increased (p=0.047) in older female mice (Fig.4d). WGA-immunofluorescence staining of histological sections of female Zeb1cKO mice at the age of 25 weeks showed a non-conformal pattern in comparison to wild type mice and quantification of the cardiomyocyte cross-sectional area showed increased cardiomyocyte size (p<0.0001) (Supp.Fig.3d) and Masson’s Trichrome staining confirmed dilation of both right and left ventricles in Zeb1cKO mice (Fig.4e).

**Figure. 4.**
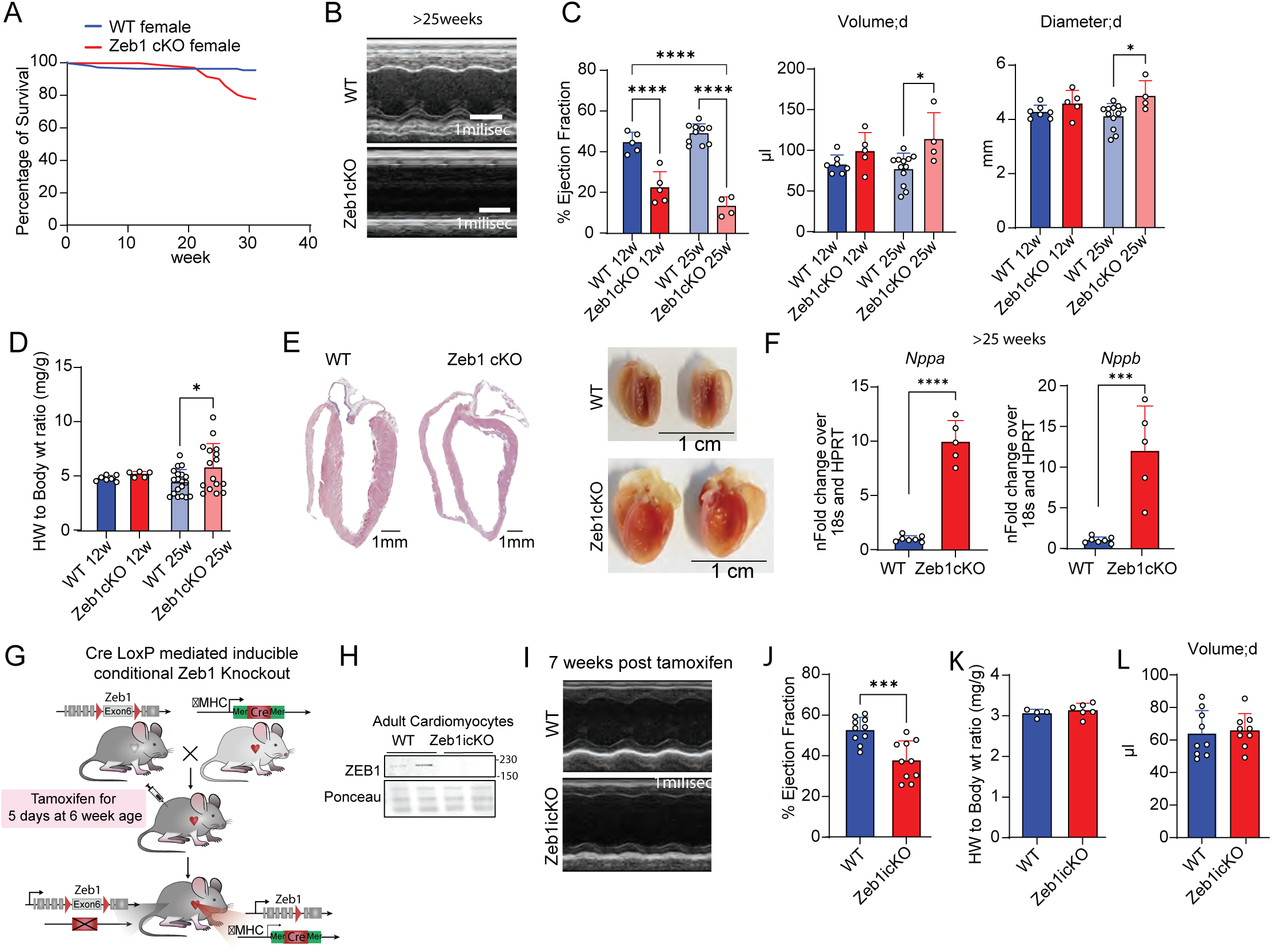
Female Zeb1 knockout mice exhibit severe cardiac dysfunction. (a) Survival rate of female Zeb1cKO mice in comparison to wild type female mice by Kaplan Meyer curve, counting the number of mice dying naturally in the animal. Zeb1cKO have a reduced survival post-20 weeks of age. (b) Representative M-mode image showing severe LV dilation of 25 weeks and older female Zeb1cKOs. (c) Age wise comparison of EF, LV volume, and diameter measured by echocardiography of female Zeb1cKO showing decreased cardiac function by 12 weeks of age and further deterioration. The 25 weeks and older mice show increased left ventricle volume and diameter (n=4-12/group). (d) Significant increase in Heart/body weight ratio by ≥25 weeks (n=4-12/group). (e) Masson’s Trichrome staining and heart images after formaldehyde fixation confirm enlargement of the heart and ventricular dilation in 25 weeks and older female Zeb1cKO mice. (f) *Nppa/Nppb* mRNA levels are elevated in Zeb1cKO left ventricle lysates, measured by RT-qPCR. (g) Schematic representation of breeding scheme for Zeb1icKO model using the Cre-loxP system with the inducible MerCreMer. (h) Confirmation of cardiomyocyte specific Zeb1 loss by western blot of isolated adult cardiomyocytes 3 weeks after tamoxifen injection. (i) M-mode image of the left ventricle contractions of female Zeb1cKOs 7 weeks post tamoxifen injection, showing dilation. (j) EF measurements taken from the echocardiography 7 weeks after tamoxifen injection showing only EF reduction in Zeb1icKO mice (n=10/group). (k) Unchanged heart/body weight ratio of female Zeb1icKO mice(n=4-6/group). (l) LV volume measurements taken from the echocardiography 7 weeks after tamoxifen injection showing no change in LV volume (n=10/group). ((Cardiomyocyte-specific Zeb1 conditional knockout: Zeb1cKO, Cardiomyocyte-specific Zeb1 inducible conditional knockout: Zeb1icKO, Ejection fraction: EF, Left ventricle: LV, Wild type: WT).

qPCR analysis showed that the transcripts of *Nppa* (p<0.0001) and *Nppb* (p=0.0003) were significantly higher in female Zeb1cKO LV lysates compared to WT (Fig.4f). We conclude that Zeb1 deletion causes dilatation, cardiac remodeling and dysfunction of the heart which is associated with a higher mortality in female mice compared to male.

However, the cardiomyocyte-specific KO model used in this study relies on the Zeb1 deletion in the late embryonic stage because of the Cre recombinase expression under the αMHC promoter, whose activity rises perinatally (Supp.Fig.3e). Zeb1 expression usually increases continuously during embryonic development of the heart, dropping slightly around birth and then peaking a second time days after birth (Supp.Fig.3e,f). During that second peak, Zeb1 was reported to be an important transcription factor for cell cycling genes in cardiomyocytes, affecting cardiomyocyte proliferation before the terminal differentiation [2].

To evaluate the time course of Zeb1 expression in our Knockout model, we performed immunofluorescence staining for Zeb1 and sarcomeric Actinin of sections of embryos at day 14 (E14) and hearts of postnatal day 2 (P2). We confirmed the knockout of Zeb1 in the P2 hearts, while in E14 hearts Zeb1 was expressed in the Zeb1cKO strain (Supp.Fig.3g). Further immunofluorescence staining for Ki67, a proliferation marker, was performed to assess the effect of Zeb1 perinatal deletion (39) in sections of E14 embryos and p2 hearts of wild type and of Zeb1cKO mice. In P2 hearts, where Zeb1cKO had already occurred, the number of Ki67-positive cells was significantly reduced (p=0.001) compared to wild type hearts (see Supplementary Fig. 3h, i, j).

Hence, the cardiac dysfunction and severe heart failure found in Zeb1cKO mice at least partially results from Zeb1 deletion in the late perinatal stage, potentially causing dysfunctional maturation or proliferation of cardiomyocytes.

### Inducible deletion of Zeb1 in mature cardiomyocytes reveals loss of cardiac function in adult heart

To further evaluate the role of Zeb1 in adult cardiomyocytes, we created an inducible cardiomyocyte specific Zeb1 conditional knockout (icKO) mouse model using a MERCreMER recombinase system regulated by the αMHC promoter (Fig.4g). Since the conditional αMHC-dependent Zeb1 KO had a stronger phenotype in female mice, we concentrated in studying the phenotype of the induced conditional Zeb1 KO in females. The knockout was confirmed in adult cardiomyocytes by western blot (p=0.037) (Fig.4h). Echocardiography showed, that seven weeks after ZEB1KO induction, cardiac function of Zeb1icKO mice was significantly decreased (p=0.0003) (Fig.4i,j), while the heart weight to body weight ratio and the LV volume remained unchanged (Fig.4k,l). There was no noticeable reduction in the left ventricular wall thickness of the female Zeb1icKO mice as compared to WT as evident from the Masson’s Trichrome stained heart sections (Supp.Fig.4b).

In summary, the depletion of ZEB1 in adult cardiomyocytes deteriorates heart function in female mice 7 weeks after the loss of Zeb1.

### Sarcomere disorganization in Zeb1-deficient cardiomyocytes

The analysis of the phenotype of Zeb1cKO mouse hearts showed cardiac functional impairment by Zeb1 deletion. To confirm structural changes in cardiomyocyte architecture that might cause cardiac dysfunction and dilation in Zeb1cKO mice, we used Transmission Electron Microscopy (TEM) on heart tissue from WT and Zeb1cKO female mice. A general disorganization and disarray of the myofibrils was directly evident in any of the TEM pictures, with a loss filament integrity and thicker Z-disks, was discovered by TEM (Fig.5a). To objectively evaluate these alterations, sarcomere dimensions were measured in electron microscopy images by Image J. In Zeb1 cKO mice sarcomer length was lengthened in comparison to wild type mice (p=0.022) (Fig.5b).

**Figure. 5.**
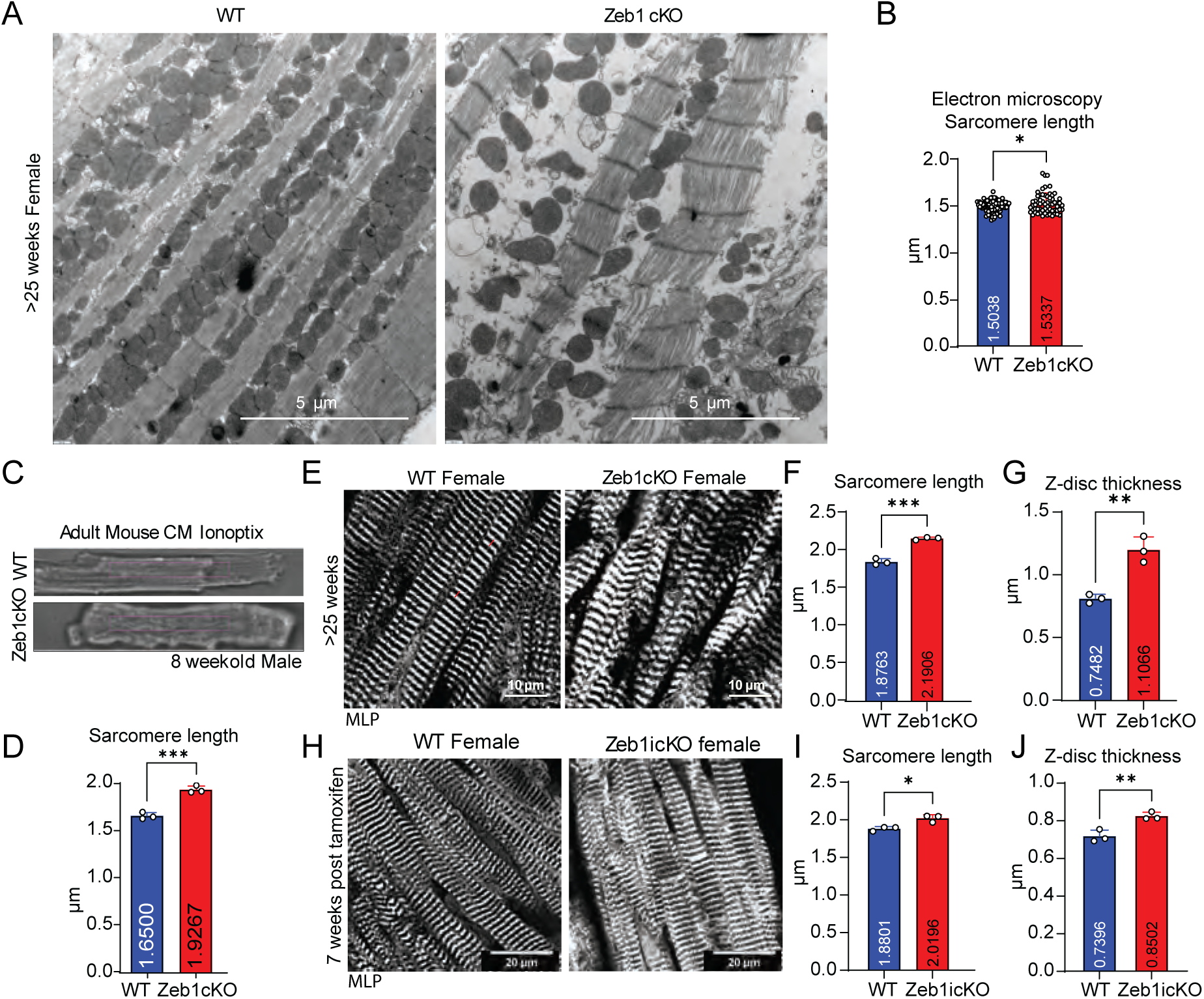
Zeb1 deletion disrupts sarcomeric structure, contributing to cardiomyocyte remodeling. (a) Representative images of TEM of 25-week-old female Zeb1cKO hearts revealed severe sarcomeric disarray and mitochondrial abnormalities. (b) Sarcomere length of hearts from 25-week-old wild type and female Zeb1cKO mice, quantified from the TEM images (n= 65-102 from 2 mice /group). (c) Representative picture of isolated ACMs from 8-week-old male wild type and Zeb1cKO mice from measurement by IonOptix. (d) Sarcomere length quantified from isolated ACMs from 8-week-old male Zeb1cKO mice, measured via IonOptix (n= 3 mice/group with 30 cells averaged). (e) Representative immage of CSRP3 immunostaining of heart sections from female wild type and Zeb1cKO heart sections. (f) Quantification of sarcomere length and (g) Z-disc thickness on immunostainings of heart sections from female wild type and of Zeb1cKO mice, revealing increased sarcomere length and Z-disc thickness in the Zeb1cKO mice. (h) Representative image of CSRP3 immunostaining of heart sections from female wild type and Zeb1icKO heart sections 7 weeks post Tamoxifen injection. (i) Quantification of sarcomere length and (j) Z-disc thickness on immunostainings of heart sections from female wild type and of Zeb1cKO mice, showing slight increase of sarcomere length and Z-disc thickness in female Zeb1icKO hearts. (Adult cardiomyocytes: ACMs, Cardiomyocytes: CM, Cardiomyocyte-specific Zeb1 conditional knockout: Zeb1cKO, Cardiomyocyte-specific Zeb1 inducible conditional knockout: Zeb1icKO, Muscle LIM-protein 2: MLP2, Transmission electron microscopy: TEM, Wild type: WT).

Given the TEM findings, we further postulated that Zeb1 deletion disrupts sarcomere structure. Using the Ionoptix system we further validated the findings of changes in the sarcomere length in isolated cardiomyocytes from WT and Zeb1cKO mice (Fig.5c). ACMs from Zeb1cKO showed a notable increase (p=0.0008) in sarcomere length compared to those from WT hearts (Fig.5c,d). Cardiomyocytes were isolated from formaldehyde-fixed frozen cardiac tissue [27] from WT and Zeb1cKO mice and the length-to-width ratio was calculated, showing a significant decrease (p<0.0001) in Zeb1cKO (Supp.Fig.5a). We performed immunostainings for the Z-disc-associated protein MLP (Muscle LIM Protein) on cardiac sections confirming visually changes in sarcomere structure (Fig.5e). Quantifications using Image J showed a notable increase (p=0.0003) in sarcomere length (Fig.5f) and Z-disc thickness (p=0.003) (Fig.5g) in Zeb1cKO mice, compared to wild type mice. Given the observed decline in cardiac function in inducible cKO mice, we hypothesized that corresponding structural alterations in the sarcomere would be detectable. Hence, we performed immunofluorescence staining’s for MLP on Zeb1icKO heart sections (Fig.5h). The sarcomere length of Zeb1icKO hearts was significantly increased (p= 0.0124) (Fig.5i), as well as the as well as Z-disc thickness (p=0.008) (Fig.5j).

In summary, the depletion of Zeb1 in perinatal cardiomyocytes results in sarcomere lengthening and Z-disc thickening, which is accompanied with cardiac dysfunction and dilation. Inducible Zeb1 KO resulted as well in sarcomere lengthening and Z-disc thickening, matching the cardiac dysfunction quantified by echocardiography.

### Bulk RNA-Seq and *de novo* motif analysis reveals that Zeb1 deficiency induces transcriptional deregulation and ectopic epithelial gene expression

As Zeb1 functions as a transcription co-regulator, bulk RNA sequencing was performed on left ventricle tissue from female WT and Zeb1cKO mice to assess gene expression differences resulting from Zeb1 deficiency. (Fig.6a). Gene Ontology (GO) term enrichment analysis identified a subset of differentially expressed genes (DEGs), predominantly associated with extracellular matrix organization and cell adhesion in the upregulated gene subset (Fig.6b), whereas the downregulated DEGs were enriched for genes involved in cardiac contraction (Supp.Fig.6a).

**Figure. 6.**
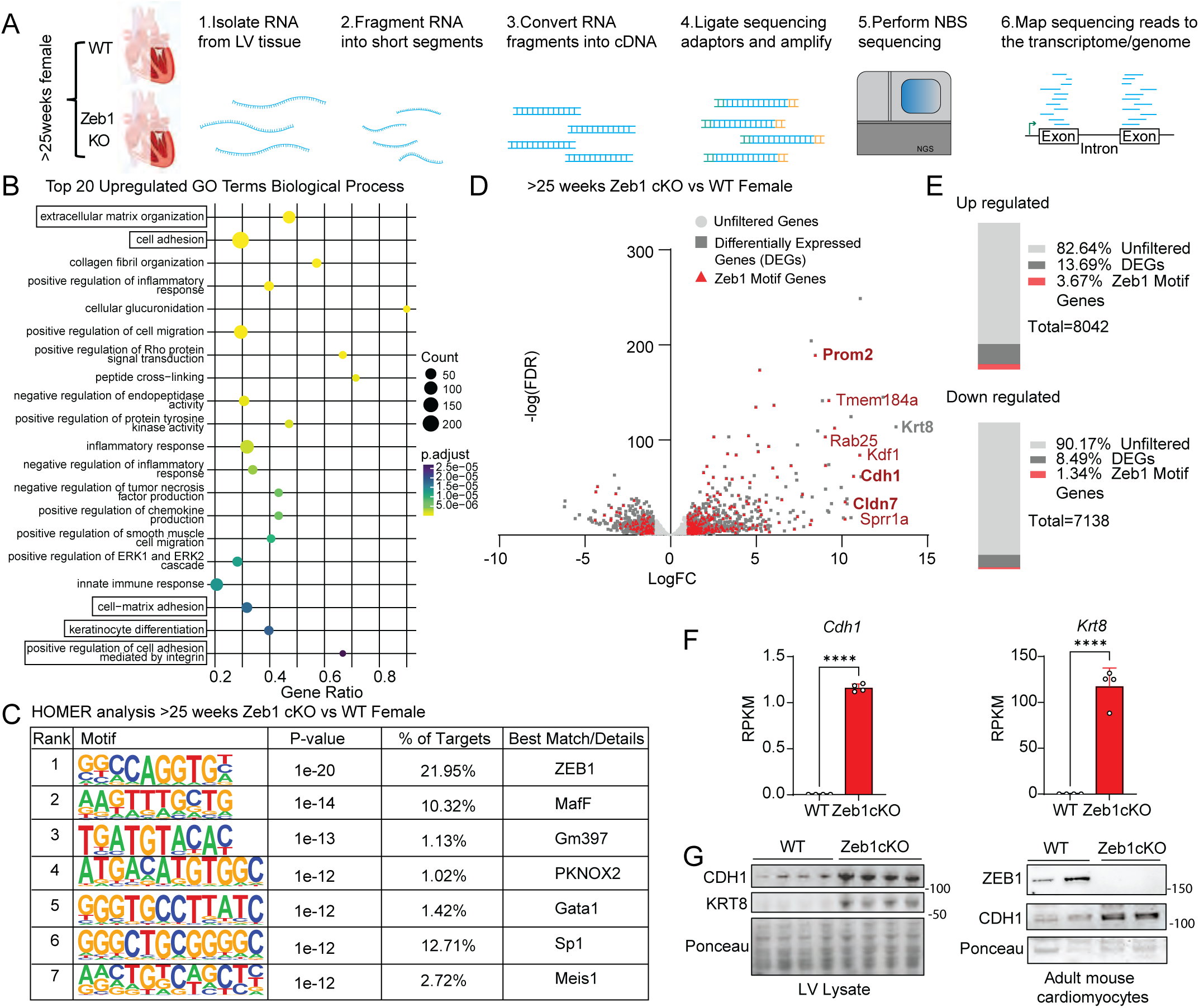
Bulk RNA-seq identifies Zeb1 as a major repressor of genes in cardiomyocytes. (a) Schematic overview of bulk RNA-Seq of left ventricle tissue from ≥25-week-old female Zeb1cKO vs. WT mice. (b) Dot-plot of top 20 GO enrichment terms for biological process of upregulated differentially expressed genes (DEGs) identified from the RNA-seq of Zeb1cKO vs WT female. Highlighted are the genes involved in ECM organization and epithelial programs. (c) De novo motif analysis (HOMER) identified Zeb1 as the top-enriched motif. (d) Volcano plot showing up and down regulated DEGs in female Zeb1cKO vs. WT mice (dark grey) and DEGs with a ZEB1 binding motif (red). (e) Percent enrichment showing Zeb1 motif genes enriched in 3.67% of upregulated vs. 1.34% of downregulated DEGs in Zeb1cKO vs. WT mice. (f) RPKM values from RNA-seq for the epithelial markers Cdh1 and Krt8, which are usually not expressed in cardiomyocytes, showing upregulation in LVs from Zeb1cKO vs. wild type. (g) Western Blotting for Cdh1 and Krt8 from lysates of LVs from Zeb1cKO vs. wild type (left) (n=4 mice /group) and for Cdh1 from isolated adult cardiomyocytes (n=2 mice /group). (Cardiomyocyte-specific Zeb1 conditional knockout: Zeb1cKO, Gene ontology: GO, Left ventricle: LV, log fold change: logFC, Wild type: WT).

To assess potential direct gene targets of the transcription factor Zeb1, which functions both as a transcriptional activator and repressor, a comprehensive *de novo* motif analysis was conducted using the HOMER v4.11 algorithm on the transcriptomes of Zeb1cKO and wild-type mice. This analysis revealed that the ZEB1 binding motif was the top-ranked motif enriched in the promoters of DEGs, suggesting a strong likelihood of direct transcriptional regulation by ZEB1 (Fig.6c). Specifically, among the 1,396 upregulated DEGs ZEB1cKO mice, 295 (3.67%) genes contained a known ZEB1 binding motif in their promoter regions. In contrast, only 96 (1.34%) out of 702 downregulated DEGs harbored a ZEB1 motif (Fig.6d,e). These findings align with the known function of ZEB1 as a transcriptional repressor of genes that need to be repressed for differentiation and maintenance of the cellular lineage.

Among the upregulated DEGs in female Zeb1cKO vs. WT mice, we found Cdh1 (p<0.0001) and Krt8 (Cytokeratin 8) (p<0.0001) as major epithelial markers (Fig.6f) among other genes like Prom2 and Cldn7 (Supp.Fig.6b). Correspondingly protein levels of these markers, which are usually not expressed in cardiomyocytes, were elevated in Zeb1cKO LV tissue lysates and adult mouse cardiomyocytes (Fig.6g).

### Integration of Zeb1 ChIP-seq and Transcriptomics Reveals Direct Regulatory Programs in Cardiomyocytes

With the aim to identify direct cardiomyocyte specific target genes, we performed Zeb1 ChIP-seq on neonatal rat ventricular cardiomyocytes for the ChIP-seq because of higher Zeb1 expression level compared to adult cardiomyocytes (Supp.Fig.7a) and a higher chromatin yield (Fig.7a).

**Figure 7.**
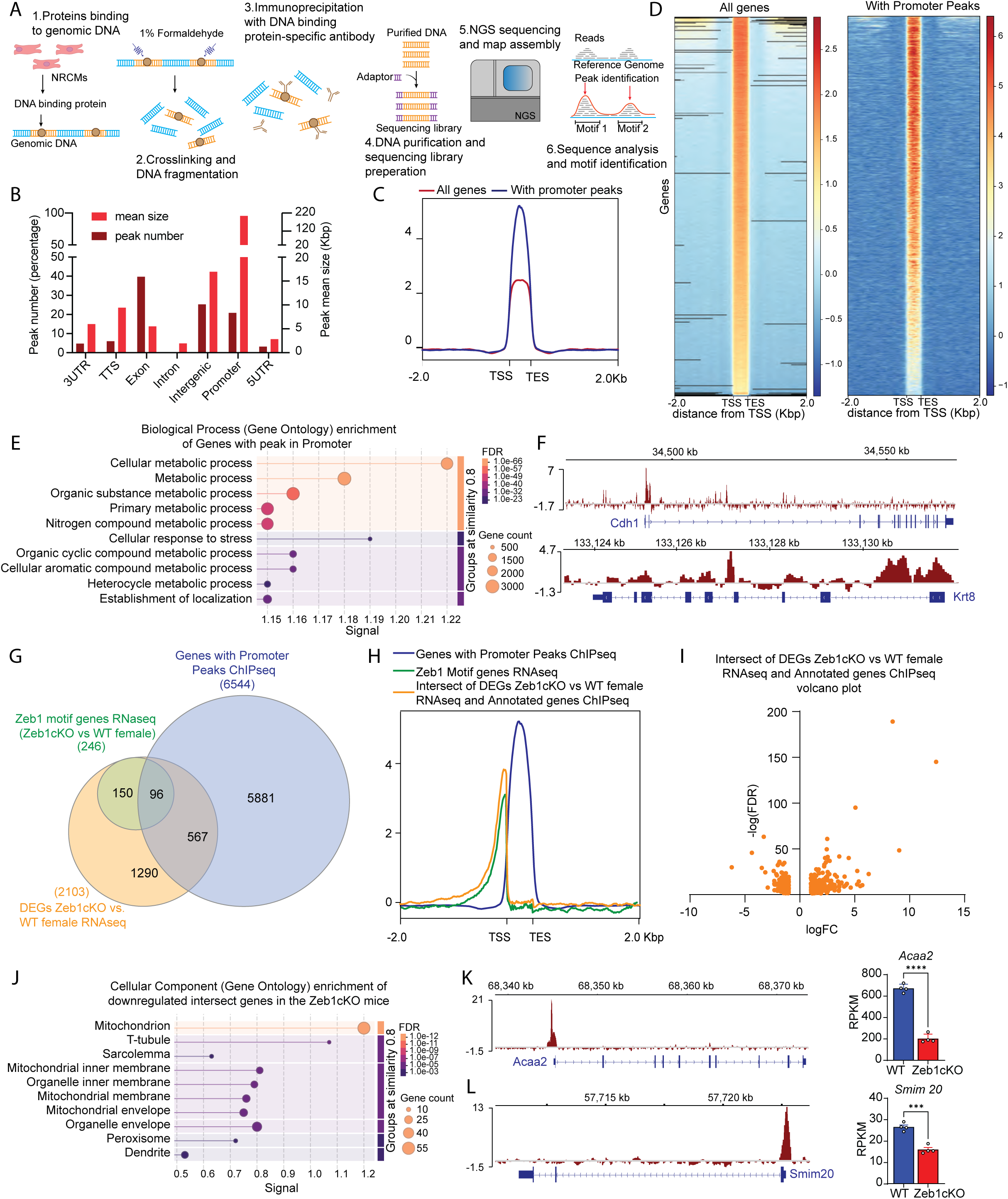
**Integration of Zeb1 ChIP-seq and Transcriptomics reveals direct activating role of Zeb1 on mitochondrial gene transcription**(a) Schematic of experimental strategy using neonatal cardiomyocytes for ChIP-seq. (b) Peak annotation showing Zeb1 enrichment in intronic (55.2%), exonic (17.8%), intergenic (11.3%), and promoter/TSS regions (12%), with promoter peaks being largest. (c, d) Heatmaps and average plot profiles of all annotated genes and genes with only promoter peaks demonstrating Zeb1 enrichment at TSSs, with sharper peaks in promoter-bound genes. (e) Gene ontology analysis for biological process of promoter-bound targets identifying enrichment in metabolic and stress-response pathways. (f) Averaged Integrated Gene view of 3 replicates of the Zeb1 ChIP-seq shows Zeb1 peaks at promoters/TSSs of Cdh1 (TSS at the left end of the gene) and Krt8 (TSS at the right end of the gene). (g) Venn diagram of promoter-peak genes (6544) and RNA-seq DEGs (2103) defines 663 high-confidence novel Zeb1 targets that are common between promoter-peak genes (6544) and RNA-seq DEGs. (h) Aggregated signal of all genes with a Zeb1 peak in the promoter area (n = 24,696) and new Zeb1 target genes identified from ChIP-seq (n = 663) show sharp TSS peaks, with the newly identified Zeb1 targets demonstrating even a sharper peak before the TSS. (i) Volcano plot of the 663 genes common between DEGs from RNA-seq of female Zeb1cKO vs WT and annotated promoter peak containing ChIP-seq genes with again less genes downregulated then upregulated in the Zeb1cKO mice. (j) GO terms of the downregulated genes described in the (i) section highlight cytoplasmic, mitochondrial, and t-tubule genes. (k) Averaged Integrated Gene view of 3 replicates of the Zeb1 ChIP-seq for two exemplary mitochondrial genes Acaa2 and (l) Smim20, showing a sharp Zeb1 binding peak at the transcription start site of Acaa2 (TSS at the left end of the gene) and Smim20 (TSS at the right end of the gene). RNA-seq reveals significant downregulation of *Acaa2* and *Smim20* in Zeb1 cKO mice (right panel of k and l). (Cardiomyocyte-specific Zeb1 conditional knockout: Zeb1cKO, Chromatin immunoprecipitation sequencing: ChIP-seq, differentially expressed genes: DEGs, Left ventricle: LV, log fold change: logFC, Transcription end site: TES, Transcription start site: TSS, Wild type: WT).

Principal component analysis (PCA) was performed on the ChIP-seq datasets to assess sample variability and overall data quality. The input samples clustered tightly together, indicating high reproducibility of the control datasets. Similarly, the three immunoprecipitated (IP) samples also grouped closely, but were clearly separated from the input samples along the first principal component, reflecting distinct enrichment patterns captured by the ChIP (Supp.Fig.7b).

Of 71,907 annotated peaks about 12,800 are localized at exonic regions (17.8%), about 40,000 (55.2%) in intronic regions and 8,146 in intergenic regions (11.3%). Approximately 8,648 (12%) of peaks from Zeb1 immunoprecipitated chromatin mapped to the Transcription Start Site (TSS) or the promoter region with the promoter peaks having the biggest mean size (Fig.7b).

To examine the genome-wide distribution of Zeb1 associated chromatin binding, we visualized input-normalized ChIP-seq read density across all annotated genes. Heatmaps and averaged density profiles revealed enrichment of Zeb1 at gene promoters, with signal peaking around the TSS. When analysis was restricted to genes with ChIP-seq peaks detected within promoter regions, a sharper enrichment profile can be observed, supporting preferential binding of Zeb1 at promoter-proximal regions (Fig.7c,d). GO enrichment analysis of these genes revealed that ZEB1 associates with genes in diverse biological processes like several metabolic processes and cellular response to stress (Fig.7e).

We further integrated Zeb1-chromatin binding with differentially expressed transcripts. Zeb1 enrichment is seen e.g. at the promoter region of Cdh1, which is a known Zeb1 target. Another example would be Krt8 with distinct Zeb1 binding in the promoter region (Fig.7f). To identify putative direct Zeb1 targets in a genome-wide manner, we compared all genes with Zeb1-binding in the promoter region to differentially expressed genes from our RNA-seq dataset of female Zeb1cKO mice and to genes containing a predicted Zeb1 motif identified by HOMER analysis within the DEGs. The Venn diagram illustrates the overlap among these datasets, highlighting the subset of genes that are both differentially expressed and promoter-bound by Zeb1 with an associated motif, representing high-confidence primary Zeb1 targets (Fig.7g). The heatmap (Supp. Fig.7d) and averaged density profiles of these 246 genes promoter-bound by Zeb1 with an associated motif, have a sharper peak before the TSS (Fig.7h).

Interestingly, we identified 567 genes that were both bound by Zeb1 at their promoters (ChIP-seq) and differentially expressed (RNA-seq), but in which HOMER motif analysis of the DEGs did not detect Zeb1 motifs. These genes may represent novel or non-canonical Zeb1 targets that were not predicted based on motif enrichment alone. However, peaks could also result from Co-immunoprecipitation of other transcription factors, without actual binding of Zeb1 to the predicted DNA binding motif. The heatmaps (Supp.Fig.7d) and averaged density profiles of these 567 genes revealed enrichment of Zeb1 at promoter region, with signal peaking before the TSS, very similar to the Zeb1 motif genes identified by RNA-seq (Fig.7h).

The expression of these 567 intersecting genes was downregulated in 171 genes whereas upregulated for 396 (Fig.7i). Gene ontology analysis of upregulated genes showed similar enrichment in cellular components of the extracellular matrix and the plasma membrane (Supp.Fig.7e). Downregulated genes with a ChIP-seq peak enriched for genes of cellular components of mitochondria, with highest count of genes and lowest FDR of p<0.00001 (Fig.7j). Inline, specific ChIP-seq analysis of the mitochondrial genes Acaa2 (Acetyl-CoA Acyltransferase 2) and Smim20 (small integral membrane protein 20) revealed sharp peaks precisely at transcription start sites. (Fig.7k). *Acaa2* and *Smim20* are significantly downregulated (p<0.001 and p<0.005) in Zeb1cKO mice, indicating that Zeb1 in this specific cases induces its expression, rather than acting as transcription repressor.

### Zeb1 regulates mitochondrial gene expression

Integrative RNA- and ChIP-seq suggest a direct Zeb1-dependent regulation of expression of mitochondrial genes. Re-evaluation our TEM pictures, we observed mitochondrial abnormalities in Zeb1cKO, which included structural disruption and a reduced number of mitochondria (Fig.8a). To confirm a direct effect of Zeb1 on mitochondria in in Zeb1cKO hearts we quantified mitochondrial DNA vs. genomic DNA in hearts, liver and skeletal muscle of Zeb1cKO and WT mice by qRT-PCR. Mitochondrial DNA vs. genomic DNA was significantly reduced in in Zeb1cKo mouse hearts (p<0.0001) in comparison to liver and skeletal muscle (Fig.8b).

**Figure 8.**
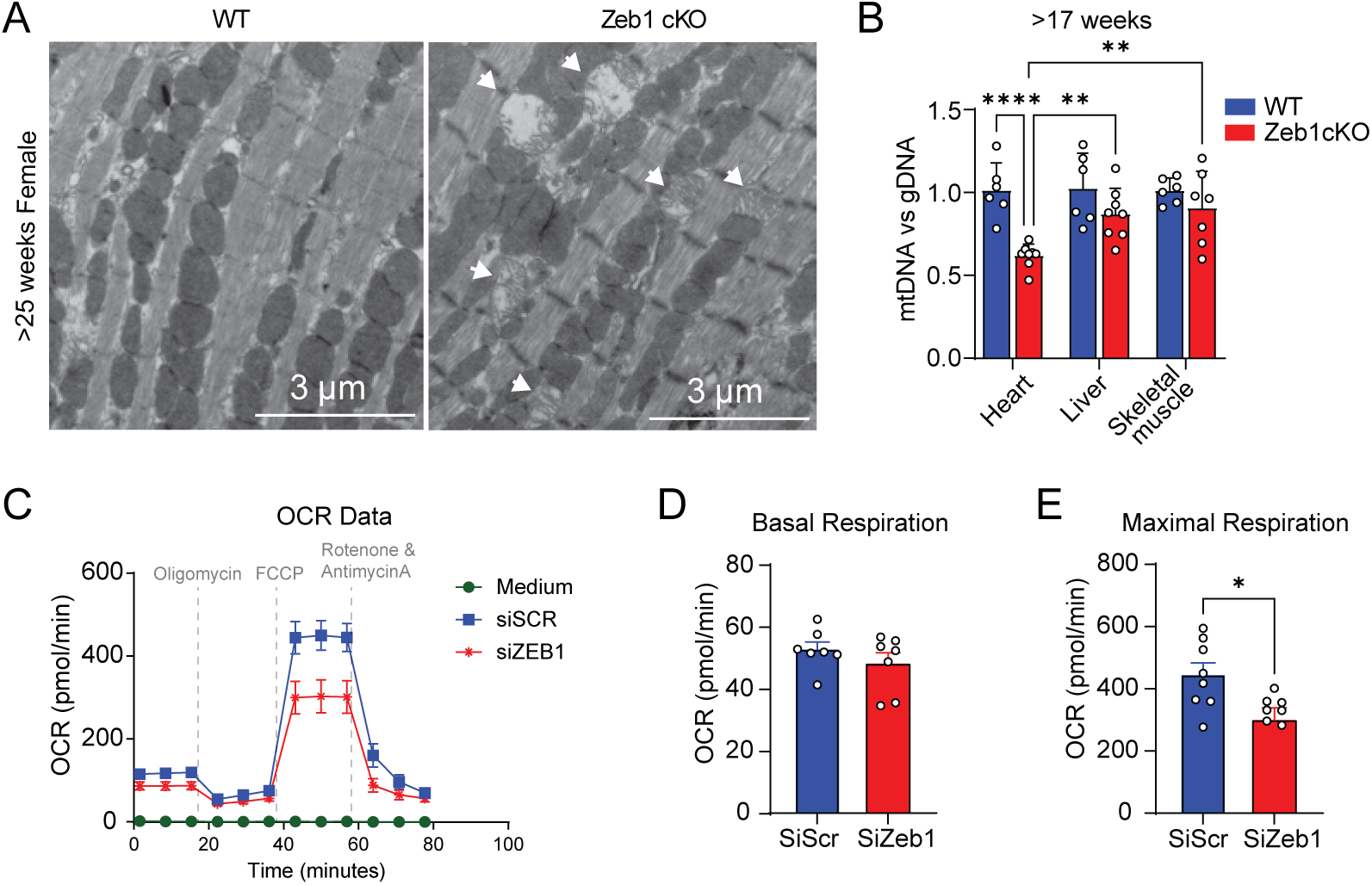
Zeb1 regulates mitochondrial integrity, abundance, and bioenergetic function. (a) Representative images of TEM of cardiac tissue from WT and Zeb1cKO hearts displaying abnormal mitochondrial morphology characterized by disrupted ultrastructure and reduced mitochondrial number in 25 weeks and older Zeb1cKO compared to WT (Arrows indicate damaged mitochondria). (b) Quantification of mitochondrial abundance of mtDNA relative to nuclear gDNA in WT and Zeb1cKO heart, liver, and skeletal muscle tissue lysates, by qRT–PCR analysis. (c) Seahorse extracellular flux analysis of OCR in NRVMs transfected with control or Zeb1-targeting siRNA. (d) Basal OCR in Zeb1-depleted cells compared to controls. (e) Maximal respiratory capacity of Zeb1 knockdown cells relative to control upon FCCP-induced uncoupling. (Cardiomyocyte-specific Zeb1 conditional knockout: Zeb1cKO, Carbonyl cyanide-4-(trifluoromethoxy)phenylhydrazone: FCCP, Genomic DNA: gDNA, mitochondrial DNA: mtDNA, Oxygen consumption rate: OCR, Transmission electron microscopy: TEM, Wild type: WT).

Since the transcriptomic data indicated downregulation of several key mitochondrial genes and the structural data mitochondrial damage, we functionally assess how the of mitochondrial genes and reduced number of mitochondria in Zeb1 depleted cardiomyocytes impact cellular bioenergetics. We performed Seahorse assays on NRCMs depleted for Zeb1 by siRNA and seahorse assay revealed that KD of Zeb1 significantly reduced mitochondrial respiration compared to controls (Fig.8c). Under basal conditions, OCR was modestly decreased in Zeb1KD cells (Fig.8d), but the impairment became significantly reduced (p=0.02) following FCCP treatment, where maximal respiration reached only ∼300 pmol/min compared to ∼450 pmol/min in controls (Fig.8e). This reduction in maximal capacity reflects a loss of spare respiratory capacity in the KD cells. These results suggest that knockdown of Zeb1 in cardiomyocytes leads to compromised oxidative phosphorylation efficiency, presumably by direct downregulation of mitochondrial components.

## Discussion

This study uncovers a previously unrecognized and critical role of the transcription factor ZEB1 in regulating cardiac structure and function. Our findings indicate that Zeb1 expression is regulated during cardiac stress and disease, and that both gain and loss of Zeb1 function led to cardiac dysfunction. We found no evidence of compensatory upregulation of Zeb2, the closest homolog of Zeb1, in either young or aged Zeb1cKO hearts. Despite sharing similar zinc finger and homeodomain structures and overlapping roles in other systems, such as during EMT and neural development, Zeb2 does not appear to compensate for the loss of Zeb1 in cardiomyocytes [17, 40].

In mice models of genetic HCM (Mybpc3 KO) mouse, we found a tendency of upregulation of ZEB1 protein levels are upregulated independently of mRNA changes, confirming a post-transcriptional regulatory mechanism as reported before [34]. Conversely, ZEB1 levels are reduced in the ICM and DCM human patient samples and a similar tendency is found in a mouse model of DCM (Rbm20 KO).

*In vitro* overexpression of Zeb1 in cardiomyocytes recapitulates key features of pathological remodeling, such as increased cardiomyocyte size and upregulation of fetal gene markers (e.g., *Nppa, Nppb*). *In vivo* overexpression of Zeb1 for 7 weeks in a cardiomyocyte-specific manner compromised cardiac function, accompanied with increases in cross-sectional cardiomyocyte areas and wall thickening, however no augmented heart weight to body weight ratio. {[13]

The cardiomyocyte-specific deletion of Zeb1, taking place in developing cardiomyocytes around the time of birth, leads to progressive cardiac dilation and dysfunction, with severe systolic impairment and chamber dilation. The decreased length to width ratio of Zeb1 cKO mice also point towards a restrictive postnatal growth phase. Our data indicate that the detrimental effects of the Zeb1 depletion are results of taking place in perinatal cardiomyocytes which are still proliferating. In Zeb1cKO mouse hearts we found less proliferative cardiomyocytes 2 days after birth in line to previous reports by Bak et al. [3]. This might also highlight a mechanism of function of Zeb1 similar to skeletal muscle, where in proliferating murine myoblasts, ZEB1 blocks myogenic genes expression. When cells exit the cell cycle, ZEB1 blockage is removed and differentiation proceeds and knockdown of ZEB1 accelerates myoblast differentiation [39].

Cardiac dysfunction is more severe in female mice, where a significant portion of Zeb1cKO females die within 30 weeks of age. Structural analysis reveals extensive sarcomeric disorganization and mitochondrial abnormalities in ZEB1-deficient hearts, indicating the importance of Zeb1 for the homeostasis of cardiomyocyte structure and function (13). Also, the deletion of Zeb1 in adult mice by induction through the MerCreMer system leads to changes int sarcomere structure and impaired function 7 weeks after the gene depletion in females. These findings highlight the importance for cardiac muscle cells, again similar to adult skeletal muscle. Siles at al. found that Zeb1 deficiency makes murine skeletal muscle more sensitive to injuries and leads to impaired regeneration of the muscle (13). Also, Zeb1 deficiency was shown to increase atrophy in a hindlimb-immobilization mouse model [31].

Mechanistically, RNA-seq analysis of Zeb1-deficient hearts discovered a prominent upregulation of extracellular matrix and cell adhesion genes, but also the re-expression of epithelial genes, like *Cdh1* and *Krt8*. This is consistent with a reversal of Zeb1’s canonical role in repressing epithelial genes during/after EMT [37]. Motif enrichment analysis further supports a direct transcriptional role for Zeb1 in maintaining cardiomyocyte identity by repressing epithelial and extracellular matrix genes. The disproportionate enrichment of Zeb1 motifs among upregulated genes confirms Zeb1’s function primarily as a transcriptional repressor in the heart.

Our integrative analysis of Zeb1 ChIP-seq and transcriptomics in cardiomyocytes demonstrates that Zeb1 preferentially binds promoter-proximal regions, where it promotes expression of genes that are components of mitochondria. The identification of high-confidence targets with both promoter binding and motif enrichment supports a direct regulatory role of Zeb1. With the help of Zeb1 ChIP-seq in NRCMs we newly identified a large set of genes that are directly regulated by Zeb1 in cardiomyocytes, that where not discovered as Zev1 targets by RNA-seq due to the lack of canonical Zeb1 motifs in the promoter region. Importantly, among these direct Zeb1 target genes we found a large set of mitochondrial genes, such as Acaa2 and Smim20, in Zeb1cKO mice. Hence, the mitochondrial loss, ultrastructural abnormalities, and decreased oxygen consumption rates are a direct effect of Zeb1 depletion and further implicate Zeb1 in maintaining the metabolic and energetic integrity of cardiomyocytes. These data might indicate that metabolic and energetic disbalance in Zeb1 cKO mice is the primary cause of cardiac dysfunction, leading to pathological remodeling.

The sex-biased phenotype in our knockout model potentially aligns with well-established mitochondrial differences between male and female hearts. A Large-scale mouse and human study has shown that females typically exhibit lower mitochondrial DNA content and reduced expression of mitochondrial genes [9], potentially making them more susceptible to the downregulation of those genes by Zeb1 depletion. Our ChIP-seq and RNA-seq data show that the Zeb1 directly regulates the expression of multiple mitochondrial genes, meaning that its loss imposes a further reduction in mitochondrial capacity in female hearts. Because females already start from a lower mitochondrial baseline, this transcriptional deficit disproportionately compromises their energetic reserve, providing a mechanistic explanation for the more severe cardiac dysfunction in female knockouts.

## Conclusion

In conclusion, our study reveals Zeb1 as a pivotal transcriptional regulator of mitochondrial genes and as repressor of genes maintaining cardiomyocyte identity. Through an integrative approach combining genetic manipulation and functional assays, we demonstrate that Zeb1 plays a role in the heart, supporting homeostasis under normal conditions while driving pathology when dysregulated.

The structural and functional impairments in Zeb1 depleted murine hearts are accompanied by a re-expression of epithelial gene expression, underscoring Zeb1’s role in preserving the mesenchymal-like phenotype of mature cardiomyocytes. Additionally, its direct importance for mitochondrial gene expression and function highlight a previously underappreciated link between EMT-associated transcriptional networks and cardiomyocyte function, positioning Zeb1 as both a guardian of cardiac homeostasis and a potential contributor to maladaptive processes.

## Abbreviation list

ZEB1: Zinc finger E-box-binding homeobox 1
TAC: Transverse Aortic Constriction
DCM: Dilated Cardiomyopathy
HCM: Hypertrophic Cardiomyopathy
NRCMs: Neonatal Rat Cardiomyocytes
PE: Phenylephrine
AAV9: Adeno-Associated Virus serotype 9
cKO: Conditional Knockout
icKO: Inducible Conditional Knockout
EF: Ejection Fraction

## Supplementary Figures

**Supplementary Figure 1.**
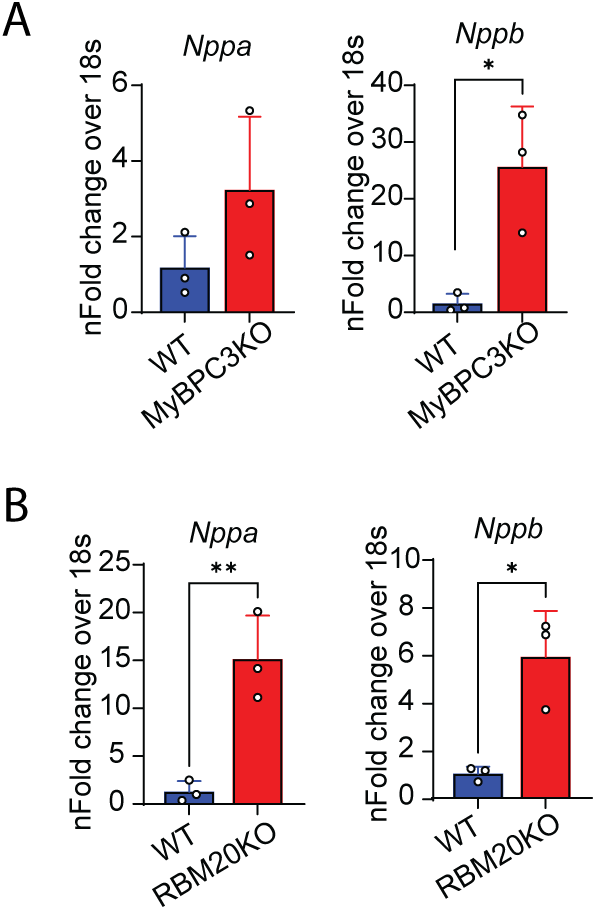
(a) Relative transcript levels of *Nppa* and *Nppb* levels are elevated in lysates of left ventricle from Mybpc3KO and (b) Rbm20KO mice, quantified by RT-qPCR.

**Supplementary Figure 2.**
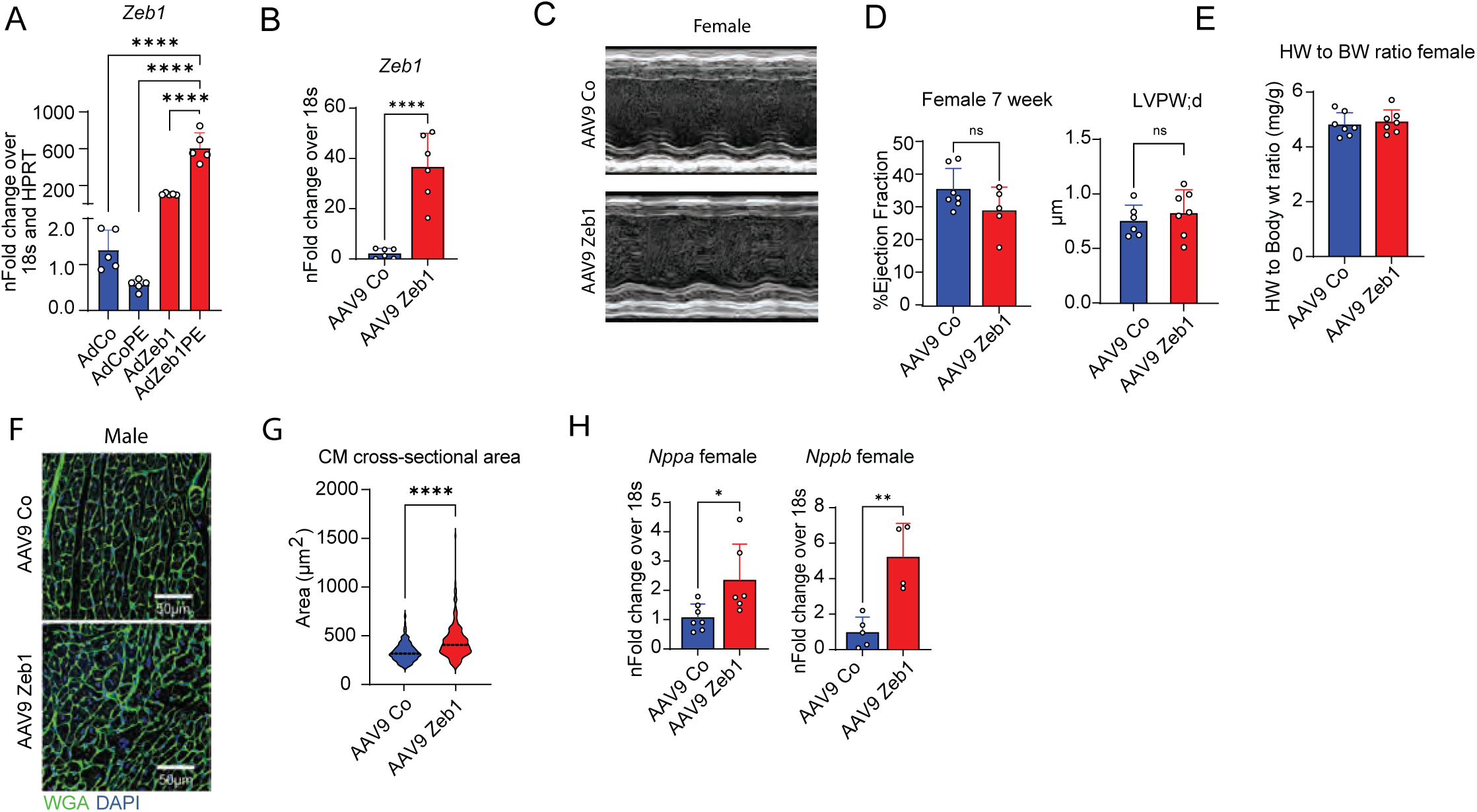
(a) Relative mRNA levels of *Nppb* are increased in Zeb1OE NRCMs as well as in PE treated NRCMs compared to control virus treated cells, quantified by RT-qPCR. (b) Relative mRNA levels of *Nppa* and *Nppb* in Zeb1OE mice are increased as compared to control virus injected mice (N=4-7/group), quantified by RT-qPCR. (c) M-mode image of echocardiography showing no significant difference between control and Zeb1OE female mice. (d) EF and LV volume measurement showing no significant change upon Zeb1OE in females (N=5-7/group). (e) Unchanged heart/body weight ratio (N=7/group). (f) WGA staining of formaldehyde fixed paraffin embedded LV tissue sections showing (g) increased cardiomyocyte cross-sectional area. The cross-sectional area was quantified and plotted (n=200 cells/group). (h) *Nppa/Nppb* mRNA expression in female mice with AAV9-mediated, cardiomyocyte-specific Zeb1 OE, quantified by RT-qPCR (n=4-8/group).

**Supplementary Figure 3.**
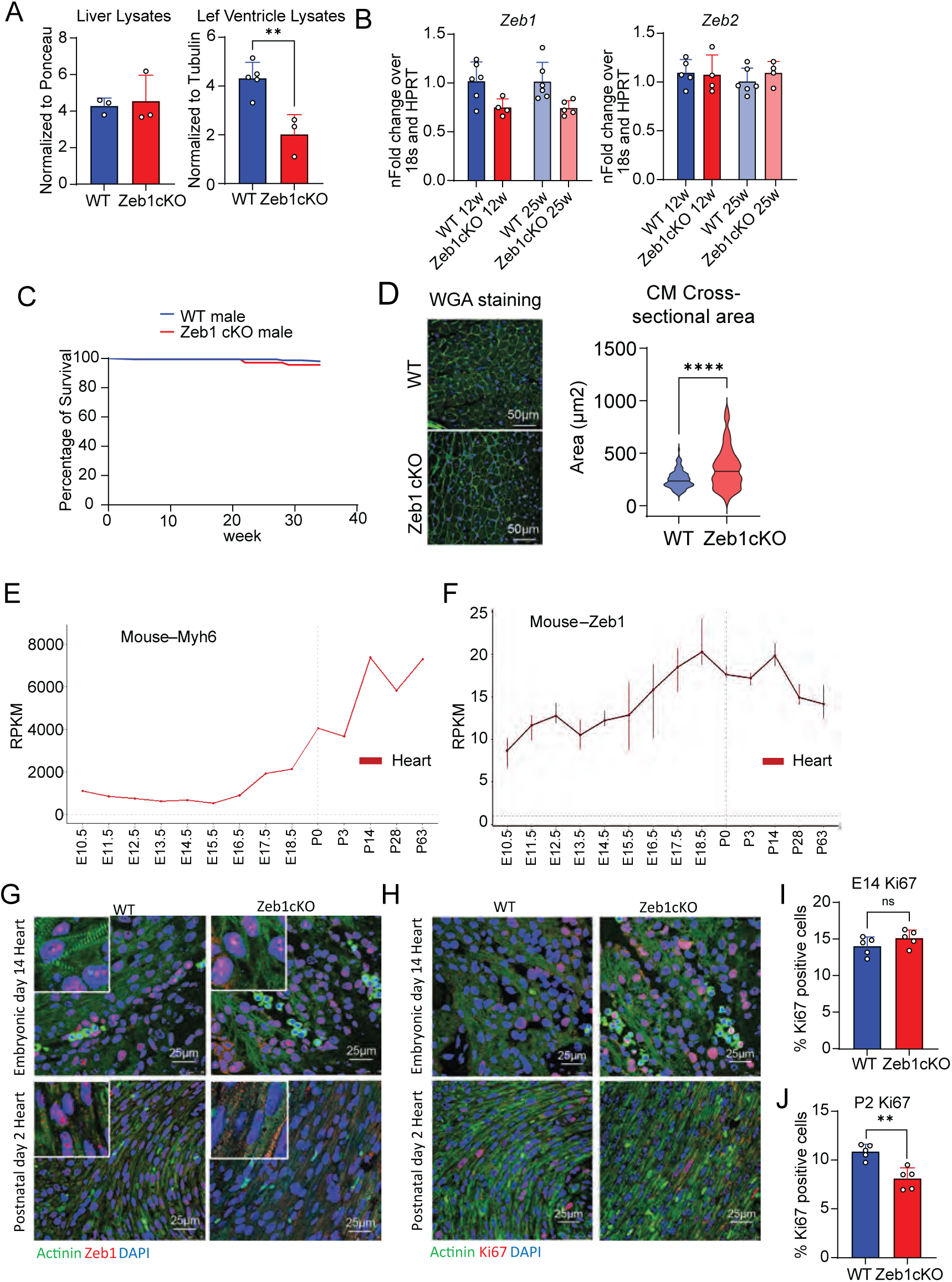
(a) Western blot quantification of Fig.3a normalized to the respective loading controls showing a significant decrease in the ZEB1 levels in LV tissue samples as compared to liver tissue samples (N=3-5/group). (b) qRT-PCR of Zeb1 and Zeb2 mRNA from the LV tissue lysates from 12 week and 25-week-old Zeb1cKO and WT mice showing reduced levels of Zeb1 mRNA but no change in Zeb2 mRNA levels (N=4-7/group). (c) Percent of survival calculated by counting the number of mice dying naturally in the animal facility showing a normal survival curve of male Zeb1 cKO mice. (d) WGA staining of formaldehyde fixed paraffin embedded LV tissue sections showing increased cardiomyocyte cross-sectional area. The cross-sectional area is quantified and plotted (n=200-250 cells/group). (e, f) Gene expression profile of Myh6 and Zeb1 in mouse generated from “Evo-devo mammalian organs” app from the Kaessmann lab. (g) Immunofluorescence staining for ZEB1 in heart sections from control and Zeb1 conditional knockout (cKO) mice at embryonic day 14 (E14) and postnatal day 2 (P2), confirming efficient Zeb1 deletion by P2 but not at E14. Nuclei were counterstained with DAPI. (h) Representative images immunofluorescence staining for Ki67 (a proliferation marker) in ventricular sections from E14 and P2 Zeb1 cKO hearts. (i, j) Quantification of Ki67-positive cardiomyocytes reveal a significant decrease in the proportion of proliferating CMs at E14 and P2 in Zeb1 cKO hearts compared to wild type (N = 5/group).

**Supplementary Figure 4.**
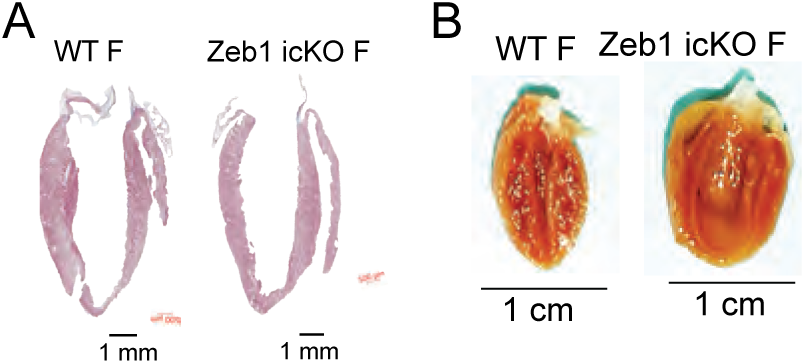
(a) Masson’s Trichrome staining and (b) heart images after formaldehyde fixation shows slight to almost no dilation in 7 weeks post tamoxifen female Zeb1icKO mice.

**Supplementary Figure 5.**
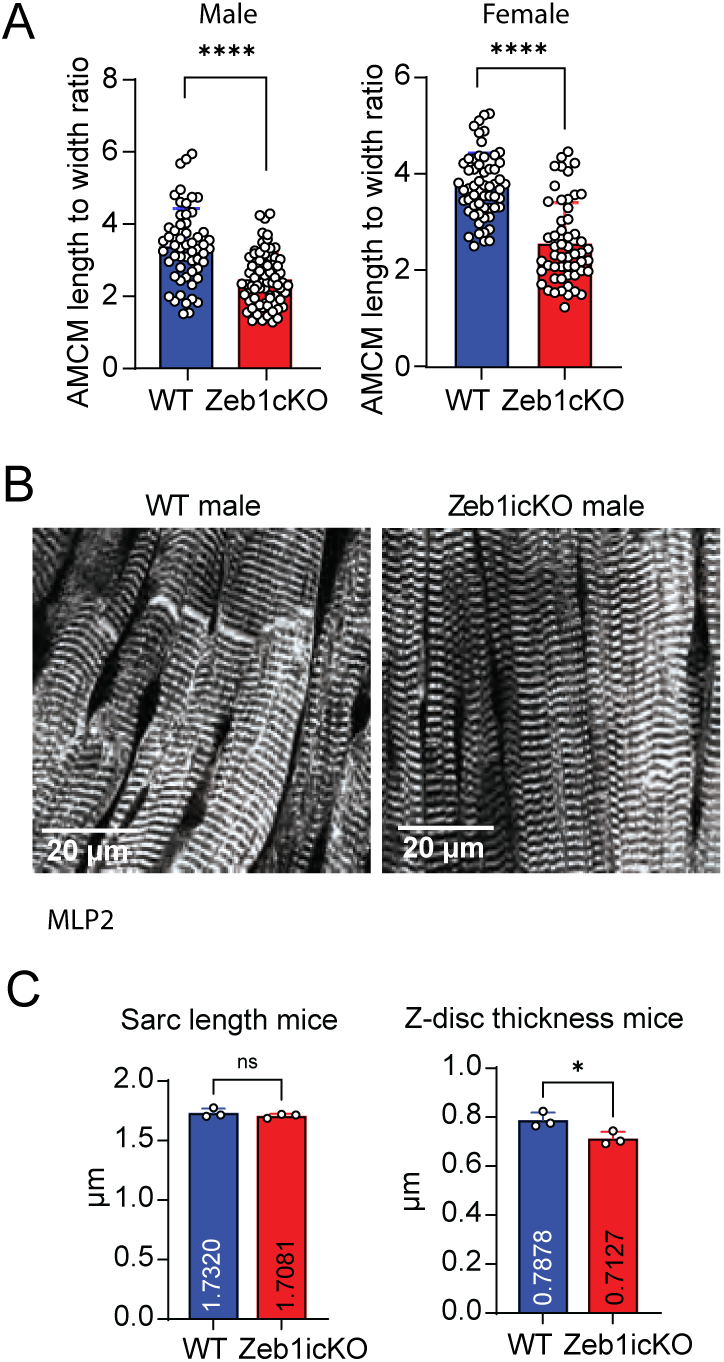
(a) Cardiomyocytes isolated from formaldehyde-fixed hearts of mice ≥25 weeks showed reduced yield in Zeb1 cKO compared to WT. Zeb1-deficient cells displayed a significantly lower length-to-width ratio, indicating altered morphology in both male and female. (b) CSRP3-based analysis of LV sections from male Zeb1 inducible conditional knockout (icKO) mice. (c) No significant change in sarcomere length and Z-disc thickness (N = 3/group).

**Supplementary Figure 6.**
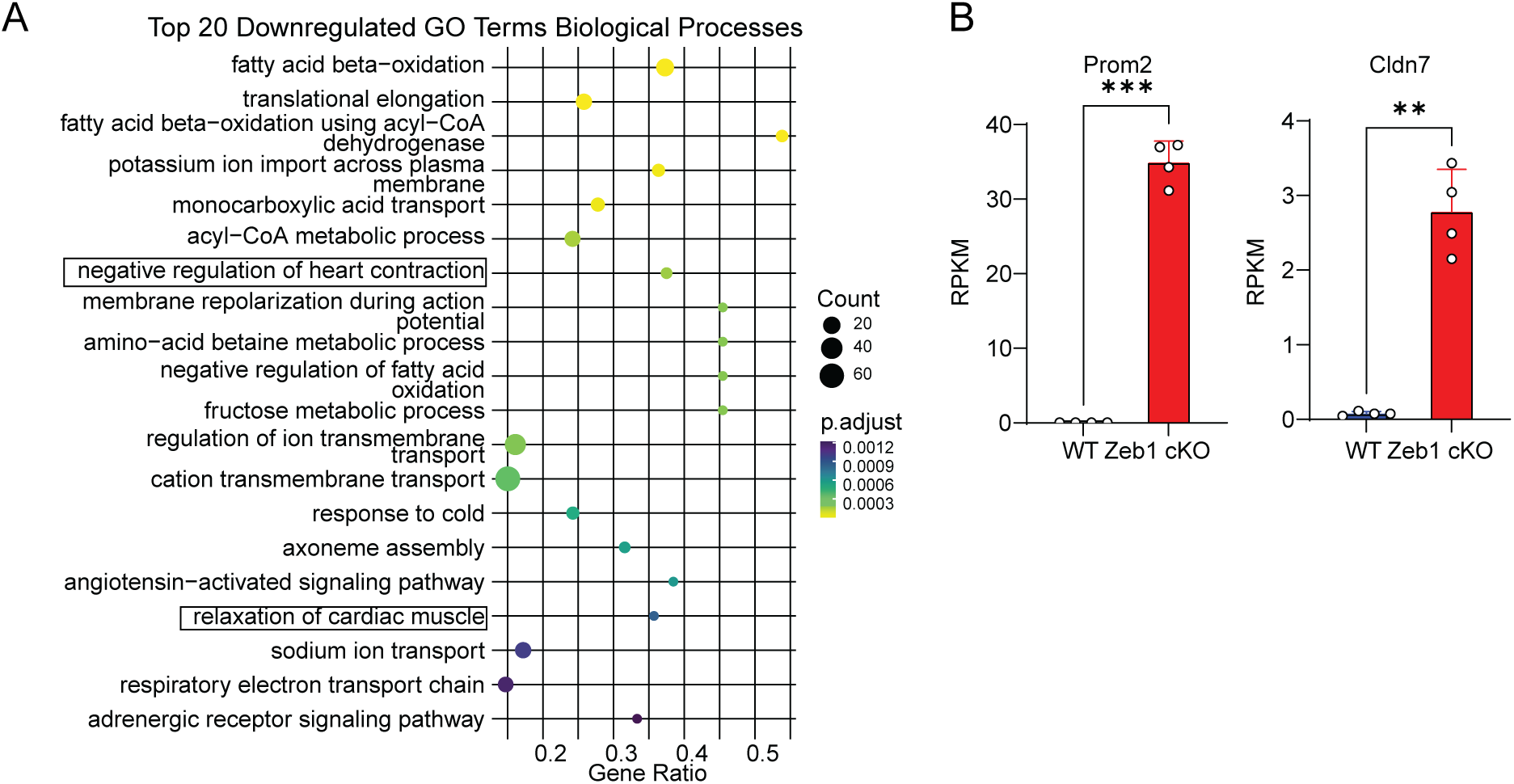
(a) Top20 downregulated Gene Ontology (GO) term enrichment analysis for biological processes of female Zeb1cKO vs WT DEGs. (b) RPKM plots of identified epithelial markers Cldn7 (Claudin 7) and Prom2 (Prominin 2) among upregulated DEGs in female Zeb1 KO mice.

**Supplementary Figure 7.** (a) qRT-PCR of Zeb1 in isolated cardiomyocytes from postnatal day 2 hearts (NCM) and 8-week-old mouse hearts (ACM). (b) Principal component analysis (PCA) of ChIP-seq datasets. PC1 on the x-axis explaining 92.4% variance and PC2 on the y-axis explaining 6.7% variance. Separated groups of input and Zeb1 samples shows good quality of ChIP-seq (N=3/group). (c) Table showing the alignment statistics of sequencing reads from input and Zeb1 immunoprecipitated (IP) chromatin. (d) Heatmaps and average plot profiles of Zeb1 motif genes identified from RNA-seq and new Zeb1 motif genes identified by ChIP-seq that overlap with RNA-seq DEGs showing enrichment before TSS, with sharper peaks in promoter regions. (e) Individual IGV tracks of all input and Zeb1 ChIP-seq replicates for Cdh1 and Krt8 showing clear peaks at the TSS and promoter regions on the Zeb1 replicates and nothing in the input replicates.

## References

1. Ackers-Johnson M, Li PY, Holmes AP, O’Brien SM, Pavlovic D, Foo RS (2016) A Simplified, Langendorff-Free Method for Concomitant Isolation of Viable Cardiac Myocytes and Nonmyocytes from the Adult Mouse Heart. Circ Res 119:909–920. doi: 10.1161/CIRCRESAHA.116.309202

2. Bak ST, Harvald EB, Ellman DG, Mathiesen SB, Chen T, Fang S, Andersen KS, Fenger CD, Burton M, Thomassen M, Andersen DC (2023) Ploidy-stratified single cardiomyocyte transcriptomics map Zinc Finger E-Box Binding Homeobox 1 to underly cardiomyocyte proliferation before birth. Basic Res Cardiol 118. doi: 10.1007/s00395-023-00979-2

3. Bak ST, Harvald EB, Ellman DG, Mathiesen SB, Chen T, Fang S, Andersen KS, Fenger CD, Burton M, Thomassen M, Andersen DC (2023) Ploidy-stratified single cardiomyocyte transcriptomics map Zinc Finger E-Box Binding Homeobox 1 to underly cardiomyocyte proliferation before birth. Basic Res Cardiol 118:1–24. doi: 10.1007/S00395-023-00979-2/FIGURES/8

4. Van Berlo JH, Molkentin JD (2014) An emerging consensus on cardiac regeneration. Nat Med 20:1386–1393. doi: 10.1038/nm.3764

5. Borlepawar A, Rangrez AY, Bernt A, Christen L, Sossalla S, Frank D, Frey N (2017) TRIM24 protein promotes and TRIM32 protein inhibits cardiomyocyte hypertrophy via regulation of dysbindin protein levels. J Biol Chem 292:10180–10196. doi: 10.1074/JBC.M116.752543

6. Brabletz S, Lasierra Losada M, Schmalhofer O, Mitschke J, Krebs A, Brabletz T, Stemmler MP (2017) Generation and characterization of mice for conditional inactivation of Zeb1. Genesis 55. doi: 10.1002/dvg.23024

7. Brabletz T, Kalluri R, Nieto MA, Weinberg RA (2018) EMT in cancer. Nat Rev Cancer 18:128–134. doi: 10.1038/nrc.2017.118

8. Buckingham M, Meilhac S, Zaffran S (2005) Building the mammalian heart from two sources of myocardial cells. Nat Rev Genet 6:826–835. doi: 10.1038/nrg1710

9. Cao Y, Vergnes L, Wang YC, Pan C, Chella Krishnan K, Moore TM, Rosa-Garrido M, Kimball TH, Zhou Z, Charugundla S, Rau CD, Seldin MM, Wang J, Wang Y, Vondriska TM, Reue K, Lusis AJ (2022) Sex differences in heart mitochondria regulate diastolic dysfunction. Nature Communications 2022 13:1 13:3850-. doi: 10.1038/s41467-022-31544-5

10. Cencioni C, Spallotta F, Savoia M, Kuenne C, Guenther S, Re A, Wingert S, Rehage M, Sürün D, Siragusa M, Smith JG, Schnütgen F, Von Melchner H, Rieger MA, Martelli F, Riccio A, Fleming I, Braun T, Zeiher AM, Farsetti A, Gaetano C (2018) Zeb1-Hdac2-eNOS circuitry identifies early cardiovascular precursors in naive mouse embryonic stem cells. Nat Commun 9. doi: 10.1038/s41467-018-03668-0

11. Chen T, Pan P, Wei W, Zhang Y, Cui G, Yu Z, Guo X (2022) Expression of Zeb1 in the differentiation of mouse embryonic stem cell. Open Life Sci 17:455–462. doi: 10.1515/biol-2022-0042

12. Chen Y, Chen L, Lun ATL, Baldoni PL, Smyth GK (2025) edgeR v4: powerful differential analysis of sequencing data with expanded functionality and improved support for small counts and larger datasets. Nucleic Acids Res 53. doi: 10.1093/nar/gkaf018

13. Dobin A, Davis CA, Schlesinger F, Drenkow J, Zaleski C, Jha S, Batut P, Chaisson M, Gingeras TR (2013) STAR: Ultrafast universal RNA-seq aligner. Bioinformatics 29:15–21. doi: 10.1093/bioinformatics/bts635

14. Ewels P, Magnusson M, Lundin S, Käller M (2016) MultiQC: Summarize analysis results for multiple tools and samples in a single report. Bioinformatics 32:3047–3048. doi: 10.1093/bioinformatics/btw354

15. Finke D, Schanze LM, Schreiter F, Kreußer MM, Katus HA, Backs J, Lehmann LH (2022) Histone deacetylase 4 deletion broadly affects cardiac epigenetic repression and regulates transcriptional susceptibility via H3K9 methylation. J Mol Cell Cardiol 162:119–129. doi: 10.1016/j.yjmcc.2021.09.001

16. Gheldof A, Berx G (2013) Chapter Fourteen - Cadherins and Epithelial-to-Mesenchymal Transition. In: van Roy F (ed) Progress in Molecular Biology and Translational Science. Academic Press, pp 317–336. doi: 10.1016/B978-0-12-394311-8.00014-5

17. Gladka MM, Kohela A, Molenaar B, Versteeg D, Kooijman L, Monshouwer-Kloots J, Kremer V, Vos HR, Huibers MMH, Haigh JJ, Huylebroeck D, Boon RA, Giacca M, van Rooij E (2021) Cardiomyocytes stimulate angiogenesis after ischemic injury in a ZEB2-dependent manner. Nat Commun 12. doi: 10.1038/s41467-020-20361-3

18. Gu Z, Eils R, Schlesner M (2016) Complex heatmaps reveal patterns and correlations in multidimensional genomic data. Bioinformatics 32:2847–2849. doi: 10.1093/bioinformatics/btw313

19. Van Den Hoogenhof MMG, Beqqali A, Amin AS, Van Der Made I, Aufiero S, Khan MAF, Schumacher CA, Jansweijer JA, Van Spaendonck-Zwarts KY, Remme CA, Backs J, Verkerk AO, Baartscheer A, Pinto YM, Creemers EE (2018) RBM20 mutations induce an arrhythmogenic dilated cardiomyopathy related to disturbed calcium handling. Circulation 138:1330–1342. doi: 10.1161/circulationaha.117.031947/suppl_file/circulationaha.117.031947_supple mental_tables.xls

20. Judge DP, Neamatalla H, Norris RA, Levine RA, Butcher JT, Vignier N, Kang KH, Nguyen Q, Bruneval P, Perier M-C, Messas E, Jeunemaitre X, De Vlaming A, Markwald R, Carrier L, Hagège AA (2015) Targeted Mybpc3 Knock-Out Mice with Cardiac Hypertrophy Exhibit Structural Mitral Valve Abnormalities. J Cardiovasc Dev Dis 2:48–65. doi: 10.3390/jcdd2020048

21. Karbassi E, Fenix A, Marchiano S, Muraoka N, Nakamura K, Yang X, Murry CE (2020) Cardiomyocyte maturation: advances in knowledge and implications for regenerative medicine. Nat Rev Cardiol 17:341–359. doi: 10.1038/s41569-019-0331-x

22. Krebs AM, Mitschke J, Losada ML, Schmalhofer O, Boerries M, Busch H, Boettcher M, Mougiakakos Di, Reichardt W, Bronsert P, Brunton VG, Pilarsky C, Winkler TH, Brabletz S, Stemmler MP, Brabletz T (2017) The EMT-activator Zeb1 is a key factor for cell plasticity and promotes metastasis in pancreatic cancer. Nat Cell Biol 19:518–529. doi: 10.1038/ncb3513

23. Lamouille S, Xu J, Derynck R (2014) Molecular mechanisms of epithelial-mesenchymal transition. Nat Rev Mol Cell Biol 15:178–196. doi: 10.1038/nrm3758

24. Langmead B, Salzberg SL (2012) Fast gapped-read alignment with Bowtie 2. Nat Methods 9:357–359. doi: 10.1038/nmeth.1923

25. Li H, Zou J, Yu XH, Ou X, Tang CK (2021) Zinc finger E-box binding homeobox 1 and atherosclerosis: New insights and therapeutic potential. J Cell Physiol 236:4216–4230. doi: 10.1002/jcp.30177

26. Liao Y, Smyth GK, Shi W (2019) The R package Rsubread is easier, faster, cheaper and better for alignment and quantification of RNA sequencing reads. Nucleic Acids Res 47. doi: 10.1093/nar/gkz114

27. Liu H, Bersell K, Kühn B (2021) Isolation and characterization of intact cardiomyocytes from frozen and fresh human myocardium and mouse hearts. In: Methods in Molecular Biology. Humana Press Inc., pp 199–210. doi: 10.1007/978-1-0716-0668-1_15

28. Liu Y, El-Naggar S, Darling DS, Higashi Y, Dean DC ZEB1 Links Epithelial-Mesenchymal Transition and Cellular Senescence. Development. 2008 Feb;135(3):579–88. doi: 10.1242/dev.007047.

29. Madany M, Thoma T, Edwards LA (2018) The Curious Case of ZEB1. Discoveries 6:e86. doi: 10.15190/d.2018.7

30. Nieto MA, Huang RYYJ, Jackson RAA, Thiery JPP (2016) EMT: 2016. Cell 166:21–45. doi: 10.1016/j.cell.2016.06.028

31. Ninfali C, Siles L, Darling DS, Postigo A (2018) Regulation of muscle atrophy-related genes by the opposing transcriptional activities of ZEB1/CtBP and FOXO3. Nucleic Acids Res 46:10697. doi: 10.1093/NAR/GKY835

32. Peinado H, Olmeda D, Cano A (2007) Snail, ZEB and bHLH factors in tumour progression: An alliance against the epithelial phenotype? Nat Rev Cancer 7:415–428. doi: 10.1038/nrc2131

33. Ramírez F, Dündar F, Diehl S, Grüning BA, Manke T (2014) DeepTools: A flexible platform for exploring deep-sequencing data. Nucleic Acids Res 42. doi: 10.1093/nar/gku365

34. Riechert E, Kmietczyk V, Stein F, Schwarzl T, Sekaran T, Jürgensen L, Kamuf-Schenk V, Varma E, Hofmann C, Rettel M, Gür K, Ölschläger J, Kühl F, Martin J, Ramirez-Pedraza M, Fernandez M, Doroudgar S, Méndez R, Katus HA, Hentze MW, Völkers M (2021) Identification of dynamic RNA-binding proteins uncovers a Cpeb4-controlled regulatory cascade during pathological cell growth of cardiomyocytes. Cell Rep 35. doi: 10.1016/j.celrep.2021.109100

35. Roehr JT, Dieterich C, Reinert K (2017) Flexbar 3.0 - SIMD and multicore parallelization. Bioinformatics 33:2941–2942. doi: 10.1093/bioinformatics/btx330

36. Sánchez-Tilló E, Lázaro A, Torrent R, Cuatrecasas M, Vaquero EC, Castells A, Engel P, Postigo A (2010) ZEB1 represses E-cadherin and induces an EMT by recruiting the SWI/SNF chromatin-remodeling protein BRG1. Oncogene 29:3490–3500. doi: 10.1038/onc.2010.102

37. Sánchez-Tilló E, Lázaro A, Torrent R, Cuatrecasas M, Vaquero EC, Castells A, Engel P, Postigo A (2010) ZEB1 represses E-cadherin and induces an EMT by recruiting the SWI/SNF chromatin-remodeling protein BRG1. Oncogene 29:3490–3500. doi: 10.1038/ONC.2010.102

38. Siles L, Ninfali C, Cortés M, Darling DS, Postigo A (2019) ZEB1 protects skeletal muscle from damage and is required for its regeneration. Nat Commun 10. doi: 10.1038/s41467-019-08983-8

39. Siles L, Sánchez-Tilló E, Lim J-W, Darling DS, Kroll KL, Postigo A (2013) ZEB1 imposes a temporary stage-dependent inhibition of muscle gene expression and differentiation via CtBP-mediated transcriptional repression. Mol Cell Biol 33:1368–1382. doi: 10.1128/MCB.01259-12

40. Skrypek N, Bruneel K, Vandewalle C, De Smedt E, Soen B, Loret N, Taminau J, Goossens S, Vandamme N, Berx G (2018) ZEB2 stably represses RAB25 expression through epigenetic regulation by SIRT1 and DNMTs during epithelial-to-mesenchymal transition. Epigenetics Chromatin 11. doi: 10.1186/s13072-018-0239-4

41. Thiery JP, Acloque H, Huang RYJ, Nieto MA (2009) Epithelial-Mesenchymal Transitions in Development and Disease. Cell 139:871–890. doi: 10.1016/j.cell.2009.11.007

42. Thiery JP, Sleeman JP (2006) Complex networks orchestrate epithelial-mesenchymal transitions. Nat Rev Mol Cell Biol 7:131–142. doi: 10.1038/nrm1835

43. Wellner U, Schubert J, Burk UC, Schmalhofer O, Zhu F, Sonntag A, Waldvogel B, Vannier C, Darling D, Hausen A Zur, Brunton VG, Morton J, Sansom O, Schüler J, Stemmler MP, Herzberger C, Hopt U, Keck T, Brabletz S, Brabletz T (2009) The EMT-activator ZEB1 promotes tumorigenicity by repressing stemness-inhibiting microRNAs. Nat Cell Biol 11:1487–1495. doi: 10.1038/ncb1998

44. Yao L, Conforti F, Hill C, Bell J, Drawater L, Li J, Liu D, Xiong H, Alzetani A, Chee SJ, Marshall BG, Fletcher S V., Hancock D, Coldwell M, Yuan X, Ottensmeier CH, Downward J, Collins JE, Ewing RM, Richeldi L, Skipp P, Jones MG, Davies DE, Wang Y (2019) Paracrine signalling during ZEB1-mediated epithelial–mesenchymal transition augments local myofibroblast differentiation in lung fibrosis. Cell Death Differ 26:943–957. doi: 10.1038/s41418-018-0175-7

45. Zhang P, Sun Y, Ma L (2015) ZEB1: At the crossroads of epithelial-mesenchymal transition, metastasis and therapy resistance. Cell Cycle 14:481–487. doi: 10.1080/15384101.2015.1006048

46. Zhao Q, Shao T, Zhu Y, Zong G, Zhang J, Tang S, Lin Y, Ma H, Jiang Z, Xu Y, Wu X, Zhang T (2023) An MRTF-A–ZEB1–IRF9 axis contributes to fibroblast–myofibroblast transition and renal fibrosis. Exp Mol Med 55:987–998. doi: 10.1038/s12276-023-00990-6

